# A genetically encoded photo-proximity labeling approach for mapping protein territories

**DOI:** 10.1101/2022.07.30.502153

**Authors:** Nir Hananya, Xuanjia Ye, Shany Koren, Tom W. Muir

## Abstract

Studying dynamic biological processes requires approaches compatible with the lifetimes of the biochemical transactions under investigation, which can be very short. We describe a genetically encoded system that allows protein interactomes to be captured using visible light. Our approach involves fusing an engineered flavoprotein to a protein of interest. Brief excitation of the fusion protein leads to local generation of reactive radical species within cell-permeable probes. When combined with quantitative proteomics, the system generates ‘snapshots’ of protein interactions with high temporal resolution. The intrinsic fluorescence of the fusion domain permits correlated imaging and proteomics analyses, a capability that is exploited in several contexts, including defining the protein clients of the major vault protein (MVP). The technology should be broadly useful in the biomedical area.

## Main Text

Protein-protein interactions (PPIs) have a pivotal role in regulating cellular function, and their dysregulation is implicated in various diseases, such as cancer and neurodegeneration.^1^ Accordingly, modulating PPIs is recognized as a promising therapeutic strategy.^2^ Methods for the characterization of PPIs are fundamental in molecular biology, and developing new tools to study protein networks is a vibrant field of research. In the last decade, proximity labeling (PL) techniques have emerged as a powerful approach for cataloging PPIs in living cells.^3^ PL utilizes engineered enzymes genetically fused to a protein of interest (POI). The PL enzyme activates an inert small-molecule substrate, generating a short-lived reactive intermediate that diffuses away from the enzyme active site to tag neighboring biomolecules covalently. The existing PL enzymes can be divided into two families – biotin-ligases and peroxidases. Biotin-ligases, such as BioID^4^ and TurboID,^5^ utilize ATP and biotin to catalyze the formation of reactive biotin-5′-AMP, which labels lysine residues on proximal proteins.^6^ Peroxidases, e.g., APEX2,^7^ catalyze the H_2_O_2_-mediated oxidation of biotin-phenol, generating a phenoxy radical which reacts with electron-rich residues, predominantly tyrosine. The biotinylation step is followed by streptavidin pull-down, allowing for protein characterization and quantification by various mass spectrometry (MS) techniques.

Compared to traditional methods for PPI characterization, such as affinity purification coupled with mass spectrometry (AP-MS), PL offers several advantages. First, the protein networks are probed in living cells and not in lysates; thus, the chances for non-physiological artifacts are reduced. Second, PL can capture transient and weak PPIs, which are often missed in AP-MS. Furthermore, it enables the characterization of insoluble protein complexes^8^ and is useful for interrogating dynamic processes, such as GPCR signaling.^9,10^ Despite these evident advantages, existing PL methods are not without limitations. For example, some biotin-ligases suffer from poor temporal resolution, as they require labeling for several hours; the engineering of TurboID has partially solved this problem by reducing the time needed for labeling down to 10 min.^5^ The peroxidase APEX2 requires much shorter incubation with H_2_O_2_ (30-60 s) to induce biotinylation; however, some applications cannot be accessed by APEX2 labeling due to the toxicity of H_2_O_2_.

The recent introduction of the photo-proximity labeling (PPL) concept offers an attractive alternative to existing PL strategies.^11,12^ This approach relies on the delivery of a chemical photocatalyst to a specific cellular site, e.g., through conjugation to an antibody. Selective targeting of the photocatalyst allows for the localized activation of the PL probe upon light irradiation, thereby tagging proximal proteins. The PPL approach has been shown to be efficient for studying cell-surface biology and cell-cell interactions,^13-15^ as well as chromatin-associated PPIs in isolated nuclei.^16^ However, the reliance of PPL strategies on peptide- or protein-based delivery modalities generally impedes application within living cells, although cell-permeable photocatalysts can still be used intracellularly in cases where prior conjugation to a targeting protein is not required.^17,18^ In addition, unlike with genetically-encoded PL systems, supplementation of an exogenous photocatalyst could result in some off-target localization, increasing the PL background.

We, therefore, envisioned a PL system that is; (i) fully genetically encoded, and (ii) triggered by visible light. Such an approach will benefit from the high spatiotemporal resolution and the benign nature of light activation while preserving the recognized advantages of genetically encoded systems, primarily the ability to perform the labeling with exquisite specificity within living cells. Here, we report such a method, termed ‘LITag’ (**L**ight-induced **I**nteractome **Tag**ging) (**Figure 1A**). We demonstrate that this tool is broadly applicable and allows snapshots of protein neighborhoods to be reliably captured following irradiation times as short as a few seconds.

**Figure 1.**
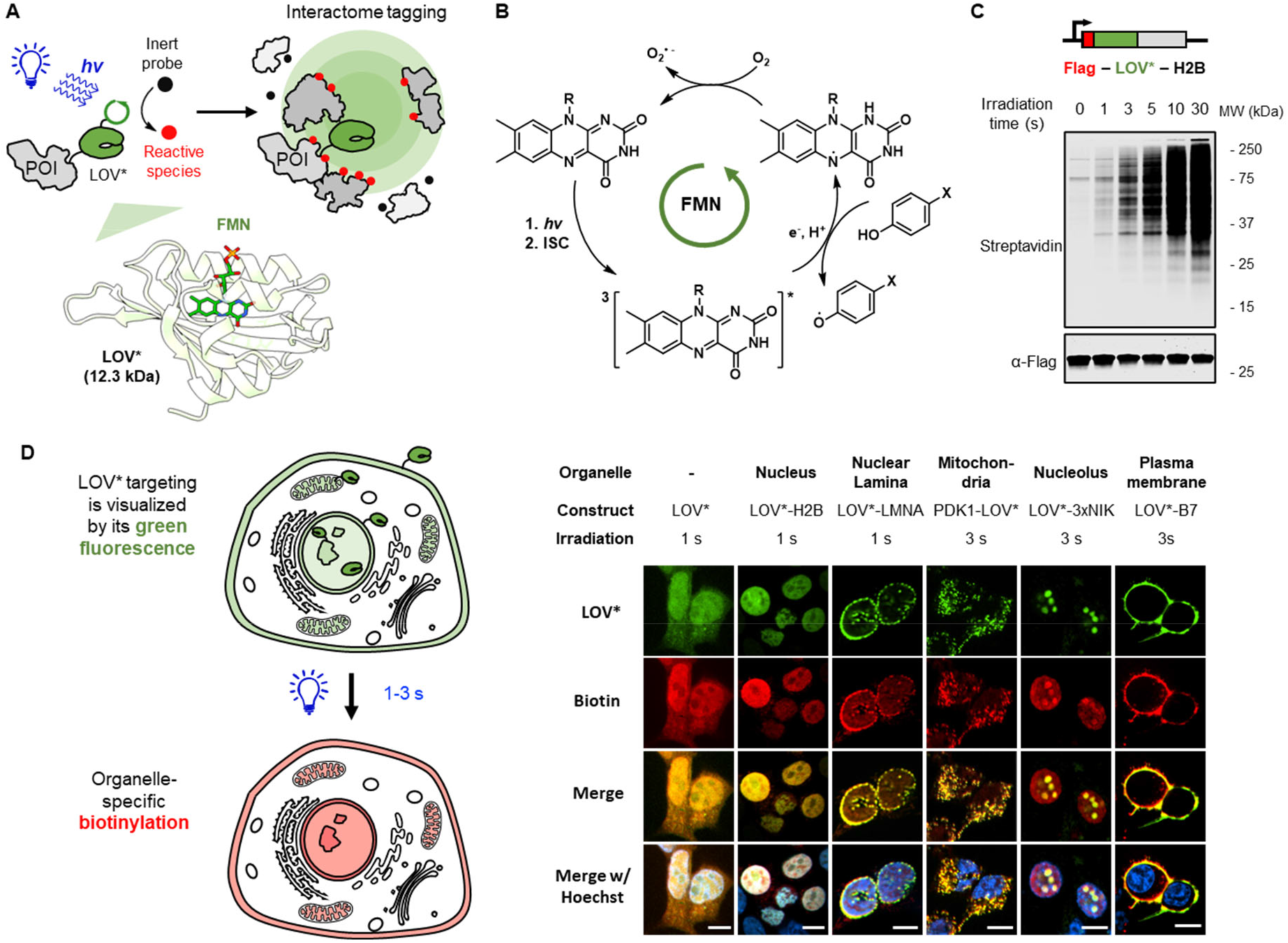
A genetically encoded approach for photo-proximity labeling.**A**. LOV*, an engineered LOV domain (PDB ID 7QF5), is fused to the POI. The FMN photocatalyst generates short-lived reactive species upon blue light irradiation, thereby tagging the POI interactome. **B**. Schematic of proposed photocatalytic cycle. **C**. Light-dependent protein biotinylation in HEK293T cells expressing a LOV* fusion. Top – map of the DNA construct used. Bottom – western blot showing protein biotinylation with different irradiation times. Cells were incubated with BP (500 µM) for 30 min before blue light irradiation. **D**. Left – LOV* is targeted to various locations based on its fusion partners and can be visualized by its intrinsic fluorescence. A short pulse of blue light is sufficient to produce localized protein biotinylation. Right – fluorescence imaging of LOV* localization and biotinylation. Live-cell LITag labeling was performed by incubating the cells with BP (BA for nucleolus labeling) and irradiating for 1-3 s. LOV* (green) was visualized by its intrinsic fluorescence (anti-FLAG antibody staining was used to visualize LOV* on the plasma membrane). Biotinylation (red) was visualized by staining with Neutravidin-Rhodamine Red-X. Hoechst 33342 (blue) is a nuclear marker. Scale bars, 10 µm.

When considering potential candidates for use in a genetically-encoded PPL system, we were drawn to the LOV (Light-Oxygen-Voltage) domains, a family of small flavin mononucleotide-(FMN-) binding proteins that have found widespread use in the optogenetics area.^19-23^ Flavin derivatives are known to participate in single electron transfer (SET) reactions upon blue-light excitation into a triplet excited state.^24-26^ Indeed, antibodies conjugated to free flavin analogs have recently been used in cell surface PPL applications using phenol-based probes.^15^ We were especially interested in engineered versions of LOV domains originally developed as singlet oxygen (^1^O_2_) sensitizers for EM imaging applications and, more recently, for labeling nucleic acids.^27-30^ Importantly, transient absorption measurements indicate that the lifetime of the FMN triplet state (t_T_) can be up to 30-fold longer in the context of the engineered LOV domains compared to the free cofactor,^31^ a feature that we anticipated would engender SET chemistry with exogenously added probes with suitable oxidation potentials (**Figure 1B**).

To test this idea, we fused a series of engineered LOV domains to histone H2B and expressed these in HEK293T cells. The cells were then treated with a biotin-phenol (BP) probe and irradiated with blue light (**Figure S1A**). Immunoblotting using a dye-labeled streptavidin revealed robust biotinylation of cellular proteins using a previously described mutant of an *Arabidopsis thaliana* phototropin LOV domain (herein referred to as LOV*, ref ^31^). Remarkably, labeling of cellular proteins could be detected with as little as 1 s irradiation when employing a suitable blue-light emitting apparatus (**Figure 1C, Figure S1B-E**). Such short irradiation pulses did not induce significant toxicity in cells expressing LOV* fusions (**Figure S1F**). Light-dependent labeling was also observed using a biotin-aniline (BA) probe, but not with aryl-azide or aryldiazirine warheads (**Figure S1G**), which is consistent with an oxidative SET pathway for probe activation (**Figure 1B**). Compared to BP, activation of BA by LOV* is more efficient, presumably because anilines have a lower oxidation potential than phenols.^32^ However, since some LOV*-independent activation of BA upon light irradiation was observed (**Figure S1G**), we chose BP as the preferred probe unless noted otherwise. Interestingly, pretreatment of cells with 1 mM H_2_O_2_ for as little as 60 s led to increased labeling efficiency (**Figure S1H**). This effect was not observed using purified proteins *in vitro* (**Figure S1J**), suggesting that the impact of peroxide in cells is indirect, potentially involving the amelioration of a redox-sensitive cellular quencher of the labeling reaction. With respect to this, we found that the *in vitro* reaction was less efficient in the presence of thiols such as glutathione (**Figure S1K**). Alternatively, peroxide treatment may affect the cellular probe concentration by modifying the plasma membrane permeability.^33^ Either way, all PPL experiments described herein were performed in the absence of added peroxide unless otherwise stated.

Next, we explored the scope of the LOV*-based PPL approach (hereafter LITag). The system was found to be compatible with a broad range of cell types (**Figure S2A**) and could be performed within a number of different cellular organelles and on the cell surface by using appropriate protein fusions (**Figure 1D, Figure S2B**,**C**). These studies highlight the potential of LITag for performing correlated cell imaging and proximity labeling. Indeed, we observed excellent spatial overlap between the intrinsic LOV* domain fluorescence for each fusion and the corresponding biotin immunofluorescence signal following BP labeling. In the course of these studies, we observed that the nucleolus could not be efficiently labeled using BP, despite clear targeting of the LITag fusion to this region (**Figure S2B**). However, this issue could be overcome by switching to the BA probe (**Figure 1D, Figure S2D**).

Encouraged by the imaging data, we next integrated LITag into a quantitative proteomics-MS workflow (**Figure 2A**). We initially focused on the mitochondria, an organelle that has been extensively studied using existing PL approaches,^5,34^ and that consequently provides an excellent opportunity to test the fidelity of our PPL approach. Efficient targeting of the LOV* domain to the mitochondrial matrix was achieved using a HEK293T cell line stably expressing a fusion with a mitochondrial targeting sequence (MTS) derived from COX4 (**Figure 2B**). These cells were treated with BP for 30 minutes before being irradiated for different times. Immunoblotting revealed labeling after a few seconds of exposure to blue light (**Figure 2C**), while immunofluorescence imaging of the irradiated cells indicated that this labeling was restricted to the mitochondria (**Figure 2B**). For quantitative proteomics-MS analysis, we integrated LITag (with 3 s irradiation) into a stable isotope labeling by amino acids in cell culture (SILAC) workflow (**Figure S3A, S3B**). This led to the identification of over 160 proteins that met our enrichment criteria (>1.5-fold change, FDR < 0.05) (**Figure 2D, S3C, D, Table S1**). Gratifyingly, every one of the enriched proteins is known to be mitochondrially localized based on human protein atlas annotation, as well as previous proteomics studies (**Figure 2E**).^35^ We also observed remarkable specificity with regard to submitochondrial localization. Since the radical intermediate is membrane-impermeant,^34^ only proteins in the matrix or the inner mitochondrial membrane (IMM) are expected to be labeled. Indeed, our list of hits does not include any proteins residing in the mitochondrial outer compartments – the intermembrane space (IMS) or the outer mitochondrial membrane (OMM) (**Figure 2E, Table S1**). The submitochondrial localization of one hit, ARL2, is uncertain;^36^ the LITag data suggest that it occupies the matrix or the IMM. In complementary studies, we showed that the MTS fusion LITag system could also be used to enrich mitochondrial RNA, in this case employing an RT-qPCR readout (**Figure 2F**).

**Figure 2.**
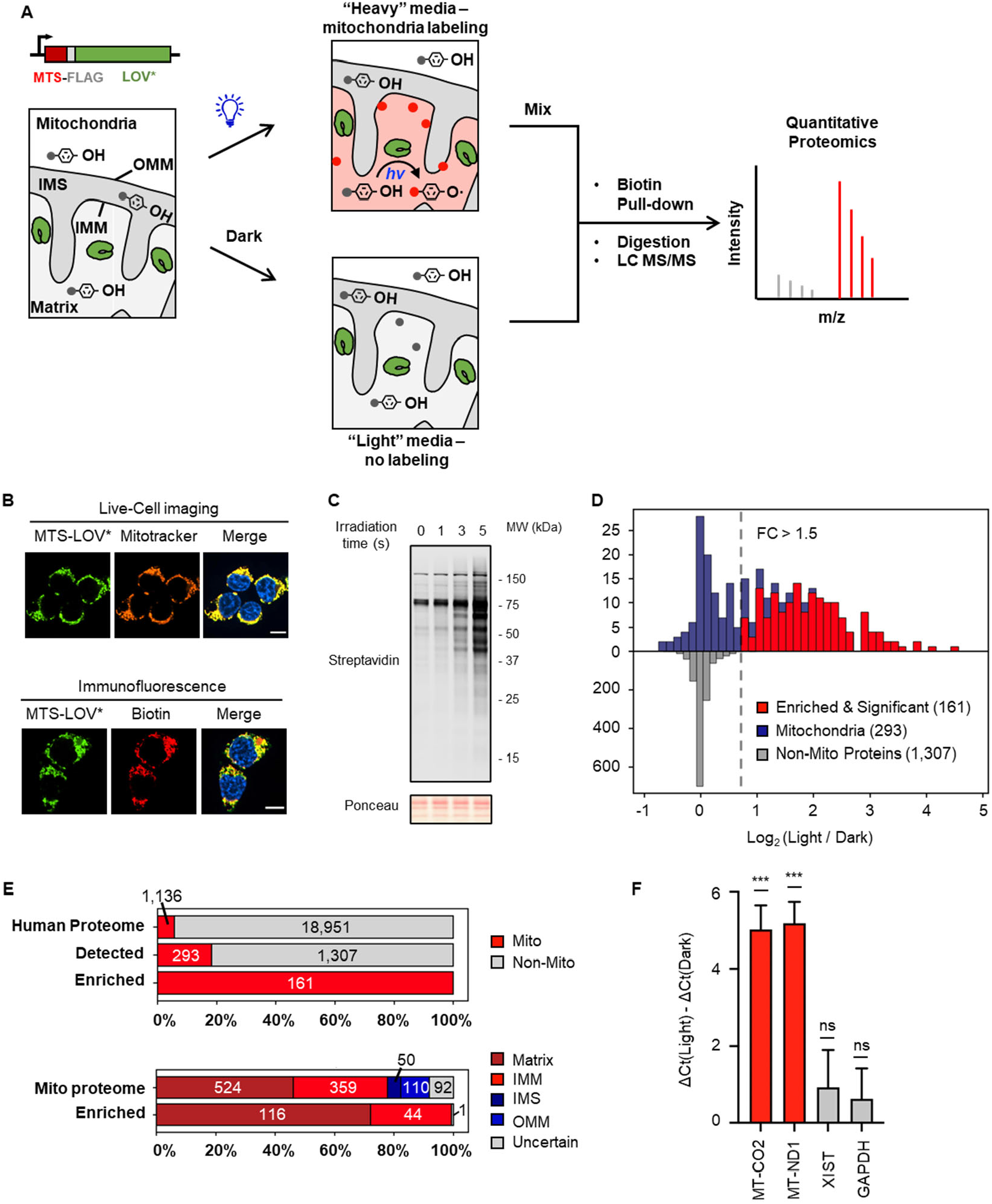
LITag labeling of the mitochondria.**A**. Workflow for LITag coupled with SILAC-based proteomics. HEK293T cells, stably expressing the MTS-LOV* fusion in the mitochondrial matrix, were cultured in either “heavy” or “light” media and incubated with BP. In the forward experiment, LITag was induced in the “heavy” cells by 3 s irradiation, while “light” cells were kept in the dark as a negative control. Biotinylated proteins were pulled-down, digested, and subjected to LC-MS/MS analysis. **B**. Top – MTS-LOV* is targeted to the mitochondria. Bottom – spatially-restricted biotinylation of the mitochondria. LOV* (green) is visualized by its fluorescence. Mitochondria (orange) are labeled with MitoTracker Deep Red FM. Biotinylation (red) is visualized by Neutravidin-Rhodamine Red-X. Hoechst 33342 (blue) is a nuclear marker. Scale bars, 10 µm. **C**. Western blot showing protein biotinylation with different irradiation times. **D**. Analysis of SILAC data from the mitochondrial matrix LITag experiment. The top histogram shows the Log_2_(FC) distribution of the mitochondrial proteins in the data set. Red bars represent significant hits (>1.5-fold change, FDR < 0.05). The bottom histogram shows the Log_2_(FC) distribution of the non-mitochondrial proteins. **E**. Analysis of specificity. Top – 1600 proteins are included in the data set, most of which (over 80%) are not mitochondrial. The labeling specificity is excellent, as only mitochondrial proteins are enriched. Bottom – labeling is restricted to the mitochondrial matrix and the IMM, with no enrichment of proteins residing in the outer compartments. **F**. LITag labeling of mitochondrial mRNAs. Biotinylated RNA molecules were pulled-down following proximity labeling, and the enrichment of 2 mitochondrial mRNAs, MT-CO2 and MT-ND1, was evaluated by RT-qPCR. No enrichment was observed for the negative controls, XIST and GAPDH. Error bars = S.D., n = 4. Data were analyzed using a one-sample *t*-test, *** p < 0.001.

As a further test of the SILAC-LITag proteomics workflow, we generated a stable U2OS cell line expressing a doxycycline-inducible LOV* fused to poly(ADP-ribose) polymerase 1 (PARP1), an essential regulator of various DNA repair pathways that catalyzes the polymerization of ADP-ribose units, attaching poly(ADP-ribose) (PAR) chains to itself and other target proteins (**Figure 3A**).^37^ The LOV* fluorescence was exploited in a live-cell imaging experiment to track the recruitment of the fusion protein to DNA damage induced by laser micro-irradiation. Indeed, PARP1-LOV* is recruited rapidly to the line of DNA damage, as expected for a functional PARP1 protein (**Figure 3B, Movie S1**).^38^ Next, the cells were treated with BP for 30 min in the presence of the damage-inducing agent H_2_O_2_. Immunoblotting revealed significant protein tagging after only 1 s of blue light irradiation (**Figure 3C**). Employing the SILAC-LITag workflow, we identified 99 hits (>1.5-fold change, FDR < 0.05) (**Figure 3D, Figure S4A**,**B, Table S2**). Most of these enriched proteins are known to be PAR binders, PARylation targets, or both (**Figure 3E**). We observed a large number of RNA-binding proteins (**Figure 3D**), consistent with the extensive involvement of RNA-processing factors in the DNA damage response.^39^ For example, the top hits on the list, FUS and EWSR1, are RNA-binding proteins that participate in DNA repair and localize to DNA breaks through PAR binding.^40,41^ We were particularly intrigued by the high enrichment of HMGB1, a multifunctional chromatin-associated protein implicated in various DNA repair pathways.^42^ HMGB1 accumulates at sites of DNA damage;^43^ however, the mechanism for this process is not yet understood. We wondered if HMGB1 recruitment is dependent upon the synthesis of PAR chains at damage sites. Indeed, laser microirradiation experiments in U2OS cells expressing an eGFP-tagged HMGB1 show that treatment with PARP1 inhibitor significantly reduces HMGB1 accumulation, strongly implicating PARylation in HMGB1 recruitment at the site of DNA damage (**Figure 3F, S4C**). Collectively, these PARP1 data indicate that the LITag approach can be used to faithfully capture a protein neighborhood during a dynamic cellular process such as the DNA damage response.

**Figure 3.**
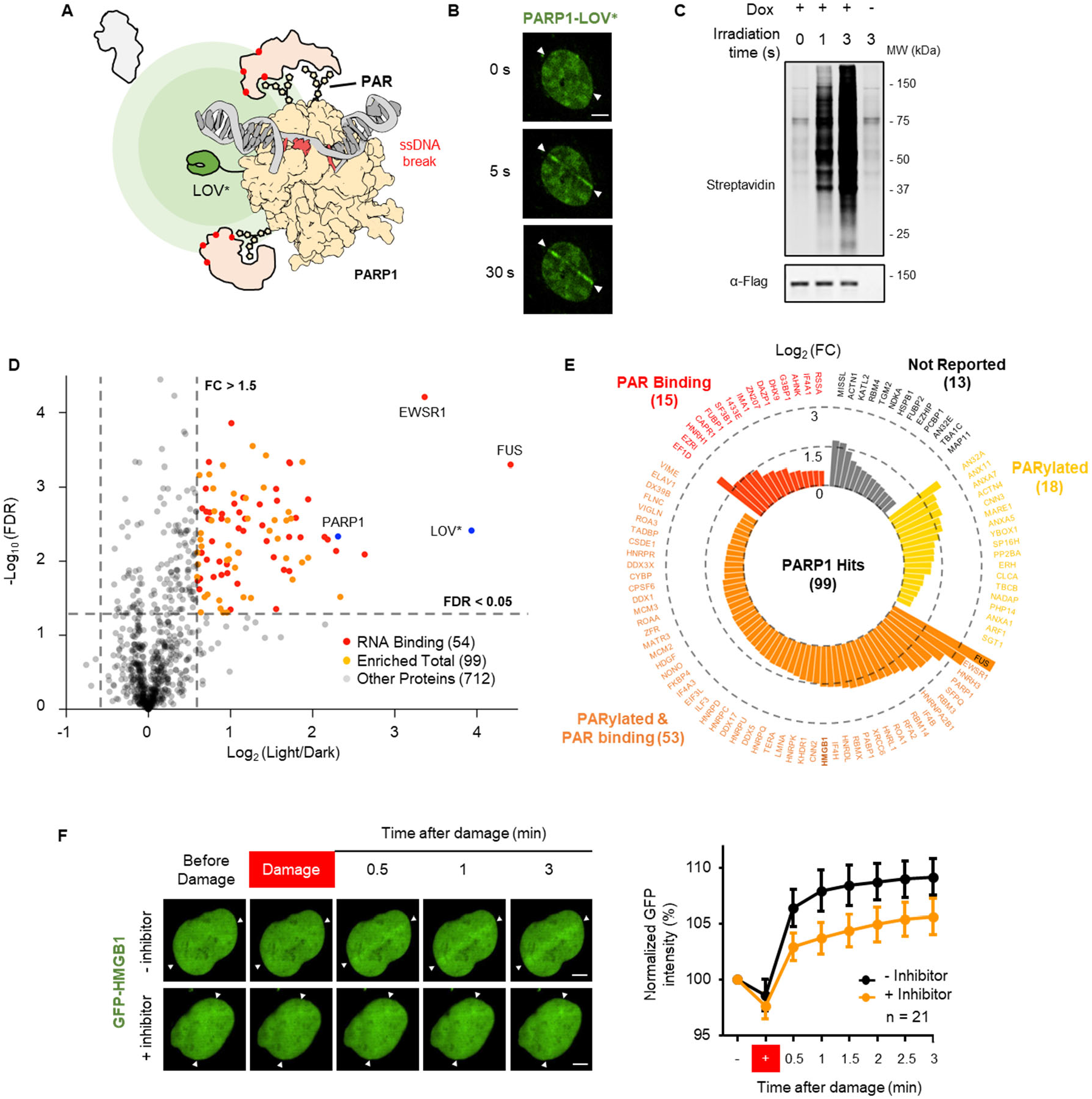
Investigation of PARP1 neighborhood following DNA damage.**A**. Treatment with genotoxic agents like H_2_O_2_ induces DNA lesions such as single-strand (ss) breaks, to which PARP1 is rapidly recruited, resulting in PARP1 activation and PAR synthesis. PAR signaling recruits various DNA repair factors, which are captured by LITag. **B**. Live-cell imaging showing the recruitment of the PARP1-LOV* fusion to the site of DNA damage induced by laser microirradiation. Fluorescence images were taken at different time points following damage induction. Scale bar, 5 µm. **C**. Western blot showing the light-dependent protein biotinylation in U2OS cells expressing PARP1-LOV*. **D**. Volcano plot displaying the SILAC-LITag data. Ninety-nine proteins were enriched (orange), among which 54 are RNA binding proteins (red). **E**. Out of 99 significant hits, 86 are known to be PAR binders, PARylation targets, or both. The enriched proteins are grouped according to these features and sorted by their log_2_ (FC), as represented by bar height. **F**. Laser microirradiation and live-cell imaging of eGFP-tagged HMGB1 in U2OS cells. Left – fluorescence images of the cells following DNA damage. Talazoparib (250 nM) was used as a PARP1 inhibitor. Scale bar, 5 µm. Right – quantification of HMGB1 accumulation at the site of DNA damage. The plus sign denotes an image taken right after microirradiation. Error bars = S.D., n = 21.

Having shown that LITag is a viable approach for PPL analyses, we next applied the strategy to characterize the interactome of the major vault protein (MVP), the primary component of the 13 MDa vault particle.^44^ MVP has been implicated in diverse cellular processes and is often associated with apoptosis resistance.^45^ However, the precise molecular mechanisms by which MVP contributes to cellular physiology remain poorly understood. In principle, characterization of the vault particle interactome by LITag could help bridge this knowledge gap. With this in mind, we fused LOV* to either the N- or C-terminus of human MVP and expressed the constructs in HeLa cells (**Figure 4A,B**). Differential centrifugation revealed that the N-terminal-tagged MVP (LOV*-MVP) is efficiently incorporated into vault particles, while the C-terminal-tagged MVP is not (**Figure 4C**). Cells expressing LOV*-MVP were incubated with biotin-containing probes and irradiated for 3 s. Robust protein tagging was observed in the presence of BA, while the use of the BP probe afforded weaker labeling in this case (**Figure S4B**); therefore, we chose BA for this LITag proteomics experiment. This led to the identification of 78 proteins as potential MVP interaction partners (>1.5-fold change, FDR < 0.05) (**Figure 4D, Figure S4C, Table S3**). PARP4, a known component of the vault particle,^46^ was highly enriched, second only to MVP itself, further validating the incorporation of LOV*-MVP into vault particles. Many of the enriched proteins are linked to nuclear transport (**Figure 4E**); this is consistent with previous literature suggesting nuclear trafficking as a major process employing the vault particle.^47,48^ Interestingly, several of the hits are chaperones or co-chaperones (**Figure 4E, Table S3**), implying a potential role for MVP in regulating protein folding. In this regard, the interaction of MVP with an HSP70 co-chaperone is implicated in apoptosis resistance.^49^ We asked whether these proteins interact with the whole vault particle or rather with the nascent MVP monomers. To answer this question, the large vault nanoparticles were isolated from soluble HeLa lysates using sucrose gradient centrifugation.^46^ Western blot analysis of the sucrose gradient fractions showed that BAG2, a nucleotide-exchange factor for the 70-kDa heat shock proteins, co-sediments with the vault particle proteins MVP and PARP4 (**Figure 4F**). This result, which was further confirmed in a non-small cell lung cancer cell line (A549, **Figure 4G**), supports the idea that MVP interacts with the folding regulators in the context of an intact vault particle. It remains to be seen whether these chaperones are actively engaged in folding client proteins while associated with the vault particle, versus being transported to where they are needed in the cell. Regardless, the MVP example indicates that the LITag approach is compatible with a large molecular particle, and, as such, may have utility in the study of analogous systems such as the assembly of viral capsids in infected cells.

**Figure 4.**
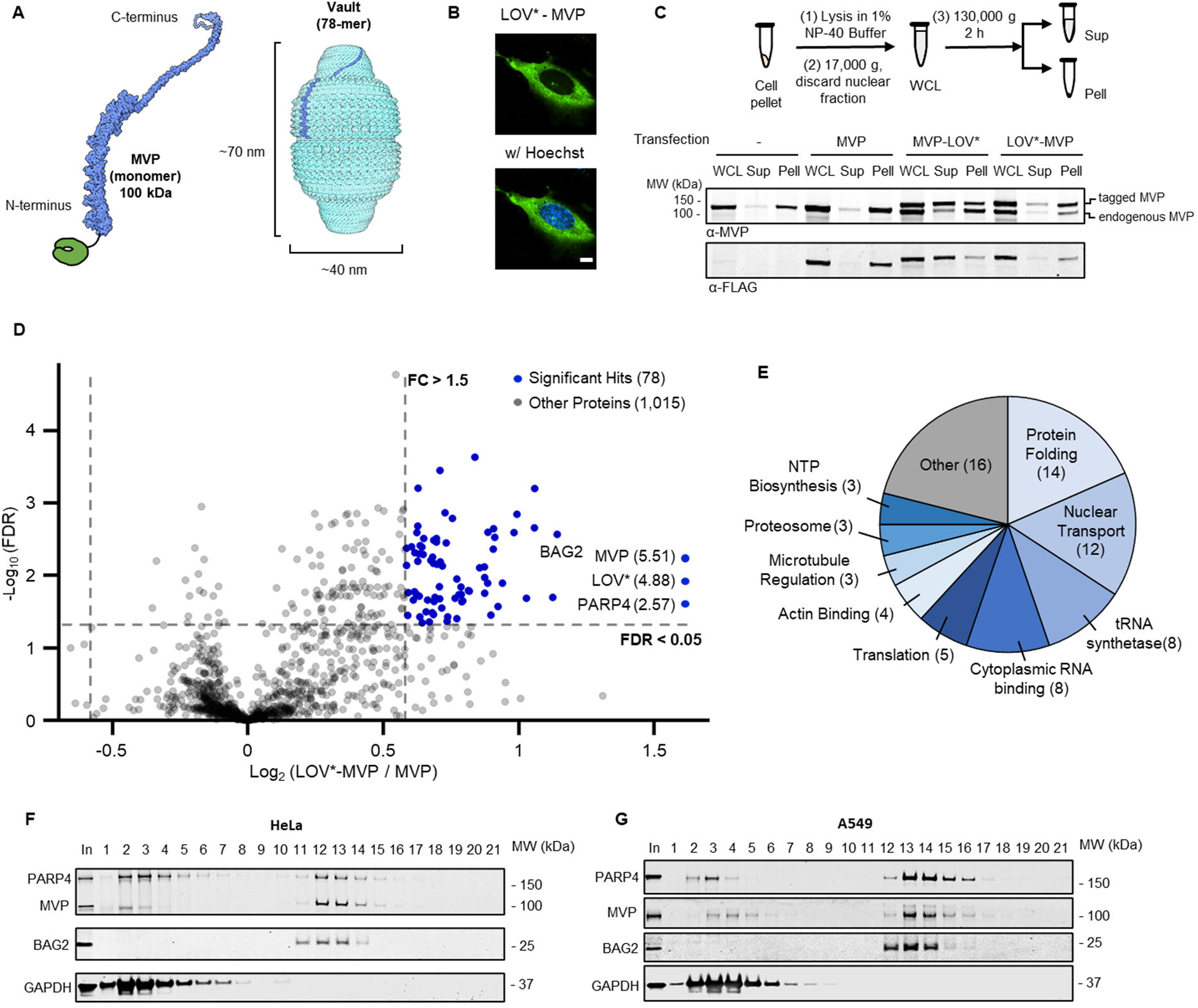
Studying the interactome of the Major Vault Protein (MVP).**A**. Right – the structure of the vault particle (PDB ID 4V60). A vault consists of 2 identical half-vaults, which align at their waists to form a barrel-like structure; each half-vault comprises 39 MVP monomers. A single monomer within the upper half-vault is highlighted in blue. Left – the small LOV* domain is fused to the N terminus of MVP. **B**. Live-cell imaging of LOV*-MVP expression in HeLa cells. LOV* (green) is visualized by its fluorescence. Hoechst 33342 (blue) is a nuclear marker. Scale bar, 5 µm. **C**. Differential centrifugation separates whole vault particles from soluble cytosolic proteins. Top – workflow: following cell lysis, the nuclear fraction is removed by centrifugation, providing a clear cytosolic fraction (WCL). The latter is subjected to ultracentrifugation; individual proteins and small complexes remain in the supernatant (Sup), while the vault particles pellet with the microsomal fraction (Pell). Bottom – western blot showing the incorporation of the exogenously-expressed MVP constructs in the vault particle. The C-terminus-tagged MVP remains largely in the supernatant, suggesting that it is not incorporated in the vault particles. On the other hand, the N-terminus-tagged MVP (LOV*-MVP) is mainly in the pellet, similarly to the endogenous MVP, implying its efficient incorporation in the vault. **D**. Volcano plot displaying the SILAC-LITag data. HeLa cells, transfected with exogenous MVP missing the LOV* domain, were used as the negative control. Seventy-eight proteins were enriched (blue) – the vault proteins MVP and PARP4 are the top hits; next is BAG2, a co-chaperone of the HSP70 proteins. MVP, LOV*, and PARP4 have been plotted arbitrarily on the right end of the x-axis. Their actual Log_2_ (FC) values are in parentheses. **E**. Enriched proteins were categorized based on their molecular function. Many hits are involved in nuclear transport and protein folding regulation. **F**,**G**. Sucrose gradient centrifugation was performed to isolate the vault particles from soluble cell lysate. Western blot analysis shows that BAG2 cosediments with the vault particles. Two different cell lines were used: **F** – HeLa; **G** – A549.

In conclusion, we have demonstrated that an engineered LOV domain can be deployed as a genetically encoded photo-proximity labeling tool. We view the LITag approach as complementary to existing PL and PLL approaches, albeit one that combines the advantages of a genetically encoded system with the spatial-temporal control afforded by an optical trigger. While the current study involved a repurposed LOV domain mutant, we imagine that further optimization of the system may be possible through directed evolution approaches, perhaps in conjunction with the use of additional cell permeable probes with suitably altered redox properties. Among the more exciting prospects that our approach offers for the future, is the application of the PPL strategy in an optically transparent model organism. Future efforts from our group will be directed along these lines.

## Acknowledgments

We thank members of the Muir laboratory for valuable discussions and comments. We thank Saw Kyin and Henry H. Shwe from the Proteomics and Mass Spectrometry Core Facility, Gary Laevsky and Sha Wang from the Confocal Imaging Facility, and Christina J. DeCoste and Katherine Rittenbach from the Flow Cytometry Resource Facility at Princeton University. We thank Dr. Steve Roffler for his helpful advice regarding the expression of LOV* on the cell surface. This work was supported by grants from the NIH [(R37-GM0968’68 and P01-CA196539 to TWM]. N.H. is a Robert Black Fellow of the Damon Runyon Cancer Research Foundation, DRG-2425-21. X.Y. is supported by a graduate fellowship from the China Scholarship Council (CSC). S.K. is supported by Human Frontier Science Program fellowship, LT000595/2020.

## Author Contributions

The project was conceived by NH, XY and TWM, with input from SK. NH, XY and SK performed all experiments. Data was analyzed by NH, XY, SK and TWM. The manuscript was written by NH, XY and TWM with input from SK.

## Data and materials availability

All data are available in the main text or the supplementary materials. The mass spectrometry proteomics data have been deposited to the ProteomeXchange Consortium via the PRIDE partner repository with the dataset identifier PXD035265. A python script for missing-data imputation of the proteomics datasets was deposited in Zenodo, DOI: 10.5281/zenodo.6883783.

## References

1. Keskin, O., Tuncbag, N. & Gursoy, A. Predicting Protein–Protein Interactions from the Molecular to the Proteome Level. Chemical Reviews 116, 4884–4909 (2016).

2. Scott, D.E., Bayly, A.R., Abell, C. & Skidmore, J. Small molecules, big targets: drug discovery faces the protein–protein interaction challenge. Nature Reviews Drug Discovery 15, 533–550 (2016).

3. Qin, W., Cho, K.F., Cavanagh, P.E. & Ting, A.Y. Deciphering molecular interactions by proximity labeling. Nature Methods 18, 133–143 (2021).

4. Roux, K.J., Kim, D.I., Raida, M. & Burke, B. A promiscuous biotin ligase fusion protein identifies proximal and interacting proteins in mammalian cells. Journal of Cell Biology 196, 801–810 (2012).

5. Branon, T.C. et al. Efficient proximity labeling in living cells and organisms with TurboID. Nature Biotechnology 36, 880–887 (2018).

6. Choi-Rhee, E., Schulman, H. & Cronan, J.E. Promiscuous protein biotinylation by Escherichia coli biotin protein ligase. Protein Science 13, 3043–3050 (2004).

7. Lam, S.S. et al. Directed evolution of APEX2 for electron microscopy and proximity labeling. Nature Methods 12, 51–54 (2015).

8. Kim Dae, I. et al. Probing nuclear pore complex architecture with proximity-dependent biotinylation. Proceedings of the National Academy of Sciences 111, E2453–E2461 (2014).

9. Lobingier, B.T. et al. An Approach to Spatiotemporally Resolve Protein Interaction Networks in Living Cells. Cell 169, 350–360.e12 (2017).

10. Paek, J. et al. Multidimensional Tracking of GPCR Signaling via Peroxidase-Catalyzed Proximity Labeling. Cell 169, 338–349.e11 (2017).

11. Geri Jacob, B. et al. Microenvironment mapping via Dexter energy transfer on immune cells. Science 367, 1091–1097 (2020).

12. Buksh, B.F. et al. μMap-Red: Proximity Labeling by Red Light Photocatalysis. Journal of the American Chemical Society 144, 6154–6162 (2022).

13. Liu, H. et al. Antigen-Specific T Cell Detection via Photocatalytic Proximity Cell Labeling (PhoXCELL). Journal of the American Chemical Society 144, 5517–5526 (2022).

14. Tay, N.E. et al. Targeted Activation in Localized Protein Environments via Deep Red Photoredox Catalysis. ChemRxiv 10.26434/chemrxiv-2021-x9bjv (2021).

15. Oslund, R.C. et al. Detection of cell–cell interactions via photocatalytic cell tagging. Nature Chemical Biology (2022).

16. Seath, C.P., Burton, A.J., MacMillan, D.W.C. & Muir, T.W. Tracking chromatin state changes using μMap photo-proximity labeling. bioRxiv, 2021.09.28.462236 (2021).

17. Trowbridge, A.D. et al. Small molecule photocatalysis enables drug target identification via energy transfer. bioRxiv, 2021.08.02.454797 (2021).

18. Tsushima, M. et al. Intracellular photocatalytic-proximity labeling for profiling protein–protein interactions in microenvironments. Chemical Communications 58, 1926–1929 (2022).

19. Pudasaini, A., El-Arab, K.K. & Zoltowski, B.D. LOV-based optogenetic devices: light-driven modules to impart photoregulated control of cellular signaling. Frontiers in Molecular Biosciences 2(2015).

20. Papanatsiou, M. et al. Optogenetic manipulation of stomatal kinetics improves carbon assimilation, water use, and growth. Science 363, 1456–1459 (2019).

21. Bracha, D. et al. Mapping Local and Global Liquid Phase Behavior in Living Cells Using Photo-Oligomerizable Seeds. Cell 175, 1467–1480.e13 (2018).

22. Strickland, D. et al. TULIPs: tunable, light-controlled interacting protein tags for cell biology. Nature Methods 9, 379–384 (2012).

23. Wu, Y.I. et al. A genetically encoded photoactivatable Rac controls the motility of living cells. Nature 461, 104–108 (2009).

24. Vaish, S.P. & Tollin, G. Flash photolysis of flavins. IV. Some properties of the lumiflavin triplet state. Journal of bioenergetics 1, 181–192 (1970).

25. Bloom, S. et al. Decarboxylative alkylation for site-selective bioconjugation of native proteins via oxidation potentials. Nature Chemistry 10, 205–211 (2018).

26. Srivastava, V., Singh, P.K., Srivastava, A. & Singh, P.P. Synthetic applications of flavin photocatalysis: a review. RSC Advances 11, 14251–14259 (2021).

27. Shu, X. et al. A Genetically Encoded Tag for Correlated Light and Electron Microscopy of Intact Cells, Tissues, and Organisms. PLOS Biology 9, e1001041 (2011).

28. Boassa, D. et al. Split-miniSOG for Spatially Detecting Intracellular Protein-Protein Interactions by Correlated Light and Electron Microscopy. Cell Chemical Biology 26, 1407–1416.e5 (2019).

29. Wang, P. et al. Mapping spatial transcriptome with light-activated proximity-dependent RNA labeling. Nature Chemical Biology 15, 1110–1119 (2019).

30. Ding, T. et al. Chromophore-Assisted Proximity Labeling of DNA Reveals Chromosomal Organization in Living Cells. Angewandte Chemie International Edition 59, 22933–22937 (2020).

31. Westberg, M., Holmegaard, L., Pimenta, F.M., Etzerodt, M. & Ogilby, P.R. Rational Design of an Efficient, Genetically Encodable, Protein-Encased Singlet Oxygen Photosensitizer. Journal of the American Chemical Society 137, 1632–1642 (2015).

32. Warren, J.J., Tronic, T.A. & Mayer, J.M. Thermochemistry of Proton-Coupled Electron Transfer Reagents and its Implications. Chemical Reviews 110, 6961–7001 (2010).

33. Funk, R.S. & Krise, J.P. Exposure of Cells to Hydrogen Peroxide Can Increase the Intracellular Accumulation of Drugs. Molecular Pharmaceutics 4, 154–159 (2007).

34. Rhee, H.-W. et al. Proteomic Mapping of Mitochondria in Living Cells via Spatially Restricted Enzymatic Tagging. Science 339, 1328–1331 (2013).

35. Rath, S. et al. MitoCarta3.0: an updated mitochondrial proteome now with sub-organelle localization and pathway annotations. Nucleic Acids Research 49, D1541–D1547 (2021).

36. Newman, L.E. et al. The ARL2 GTPase Is Required for Mitochondrial Morphology, Motility, and Maintenance of ATP Levels. PLOS ONE 9, e99270 (2014).

37. Ray Chaudhuri, A. & Nussenzweig, A. The multifaceted roles of PARP1 in DNA repair and chromatin remodelling. Nature Reviews Molecular Cell Biology 18, 610–621 (2017).

38. Haince, J.-F. et al. PARP1-dependent Kinetics of Recruitment of MRE11 and NBS1 Proteins to Multiple DNA Damage Sites*. Journal of Biological Chemistry 283, 1197–1208 (2008).

39. Mikolaskova, B. et al. Maintenance of genome stability: the unifying role of interconnections between the DNA damage response and RNA-processing pathways. Current Genetics 64, 971–983 (2018).

40. Mastrocola, A.S., Kim, S.H., Trinh, A.T., Rodenkirch, L.A. & Tibbetts, R.S. The RNA-binding Protein Fused in Sarcoma (FUS) Functions Downstream of Poly(ADP-ribose) Polymerase (PARP) in Response to DNA Damage. Journal of Biological Chemistry 288, 24731–24741 (2013).

41. Lee, S.-g. et al. Ewing sarcoma protein promotes dissociation of poly(ADP-ribose) polymerase 1 from chromatin. EMBO reports 21, e48676 (2020).

42. Lange, S.S. & Vasquez, K.M. HMGB1: The jack-of-all-trades protein is a master DNA repair mechanic. Molecular Carcinogenesis 48, 571–580 (2009).

43. Prasad, R. et al. HMGB1 Is a Cofactor in Mammalian Base Excision Repair. Molecular Cell 27, 829–841 (2007).

44. Tanaka, H. et al. The Structure of Rat Liver Vault at 3.5 Angstrom Resolution. Science 323, 384–388 (2009).

45. Frascotti, G. et al. The Vault Nanoparticle: A Gigantic Ribonucleoprotein Assembly Involved in Diverse Physiological and Pathological Phenomena and an Ideal Nanovector for Drug Delivery and Therapy. Cancers 13(2021).

46. Kickhoefer, V.A. et al. The 193-Kd Vault Protein, Vparp, Is a Novel Poly(Adp-Ribose) Polymerase. Journal of Cell Biology 146, 917–928 (1999).

47. Chugani, D.C., Rome, L.H. & Kedersha, N.L. Evidence that vault ribonucleoprotein particles localize to the nuclear pore complex. Journal of Cell Science 106, 23–29 (1993).

48. Minaguchi, T., Waite, K.A. & Eng, C. Nuclear Localization of PTEN Is Regulated by Ca2+ through a Tyrosil Phosphorylation–Independent Conformational Modification in Major Vault Protein. Cancer Research 66, 11677–11682 (2006).

49. Pasillas, M.P. et al. Proteomic Analysis Reveals a Role for Bcl2-associated Athanogene 3 and Major Vault Protein in Resistance to Apoptosis in Senescent Cells by Regulating ERK1/2 Activation. Molecular & Cellular Proteomics 14, 1–14 (2015).

