## Supplementary Material for "A genetically encoded photo-proximity labeling approach for mapping protein territories"


**This PDF file includes:**


Supplementary Figures S1-S5 ........ S2

Captions for Tables S1-S3 ........ S10

Captions for Movie S1 ........ S10

Materials and Methods ........ S11

DNA constructs ........ S20

Recombinant LOV* ........ S36

References ........ S37


**Other Supplementary Materials for this manuscript include the following:**

Tables S1-S3

Movie S1

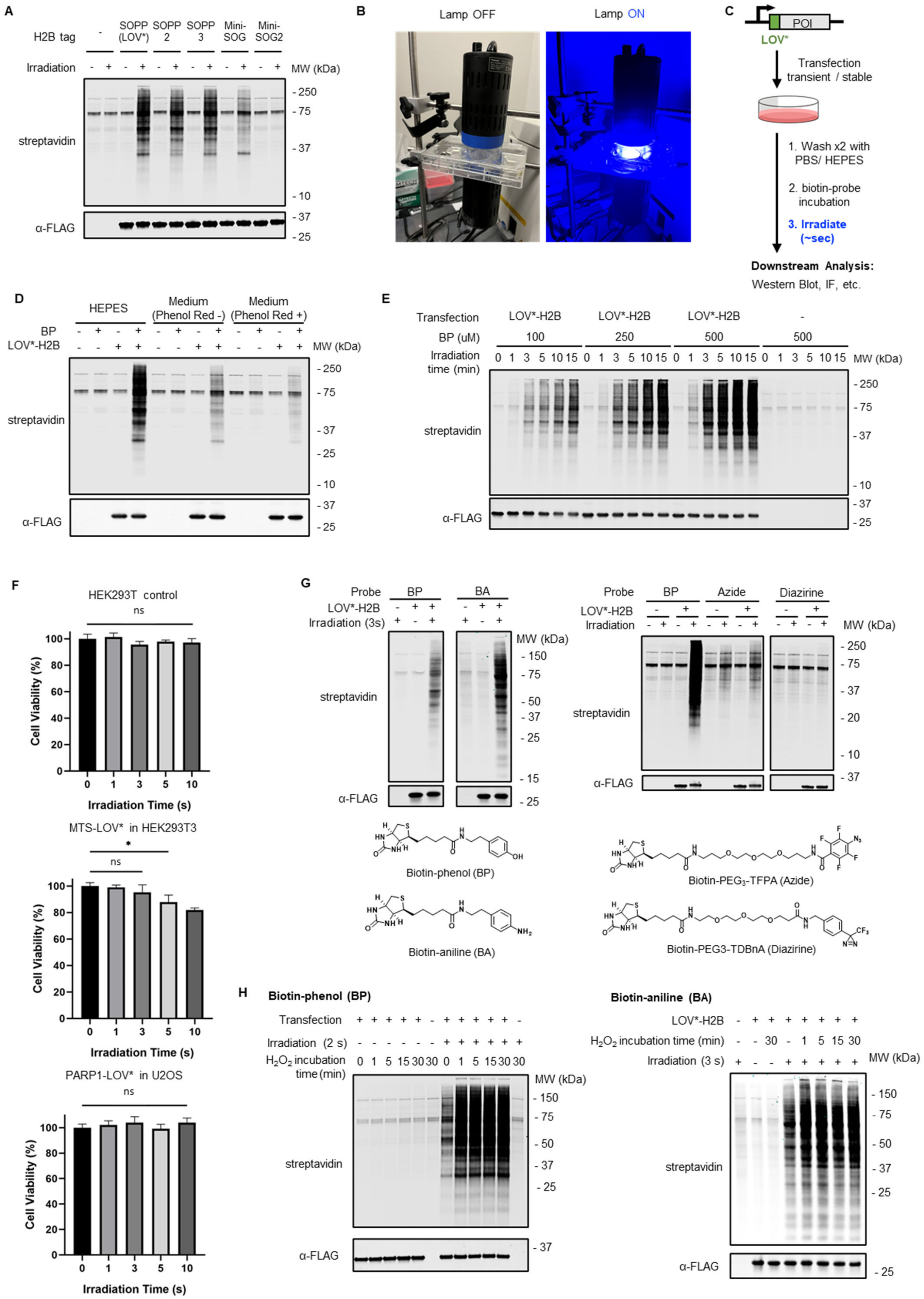
A
H2B tag
-
SOPP (LOV*)
SOPP 2
SOPP 3
Mini-SOG
Mini-SOG2
Irradiation
MW (kDa)
streptavidin
α-FLAG
B
Lamp OFF
Lamp ON
C
POI
LOV*
Transfection transient / stable
1. Wash x2 with PBS/ HEPES
2. biotin-probe incubation
3. Irradiate (~sec)
Downstream Analysis:
Western Blot, IF, etc.
D
HEPES
Medium (Phenol Red -)
Medium (Phenol Red +)
BP
LOV*-H2B
E
Transfection
BP (uM)
Irradiation time (min)
F
HEK293T control
MTS-LOV* in HEK293T3
PARP1-LOV* in U2OS
Cell Viability (%)
Irradiation Time (s)
ns
*
G
Probe
BP
BA
Azide
Diazirine
Irradiation (3s)
Biotin-phenol (BP)
Biotin-aniline (BA)
Biotin-PEG3-TFPA (Azide)
Biotin-PEG3-TDBnA (Diazirine)
H
Biotin-phenol (BP)
Biotin-aniline (BA)
Irradiation (2 s)
H2O2 incubation time (min)
Irradiation (3 s)

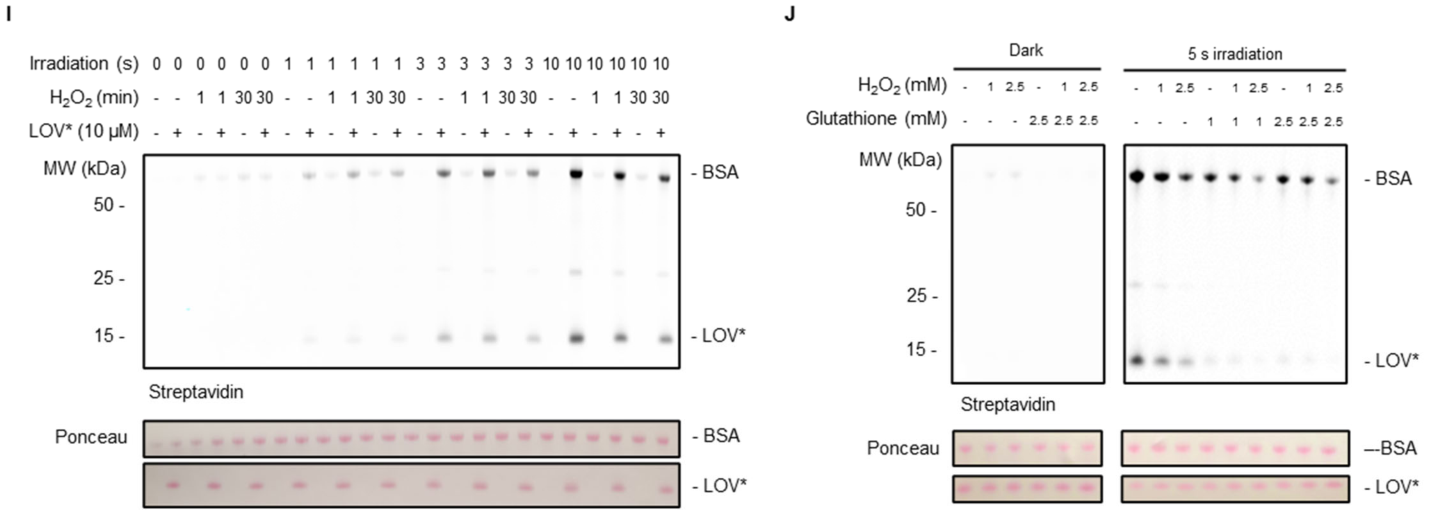


**Figure S1.** LITag method inception and optimization. **A.** Testing of engineered LOV domains for photo-proximity labeling. HEK293T cells were transfected with plasmids encoding for FLAG-tagged LOV domains fused to the N-terminus of H2B. Following incubation with biotin-phenol (BP, 500 µM), cells were irradiated with blue light, and protein biotinylation was detected by western blot using IRDye 800CW-labeled streptavidin. Western blot against FLAG was used as a loading control. MiniSOG is the archetype LOV domain engineered from the LOV2 domain of *Arabidopsis thaliana* phototropin 2 for singlet oxygen ($^1O_2$) production.[1] SOPP, SOPP2, and SOPP3 are variants of miniSOG with extended triplet state lifetimes.[2,3] MiniSOG2 was independently derived from miniSOG through directed evolution.[4] SOPP, SOPP2 and SOPP3 exhibit comparable BP activation. Since SOPP produces less $^1O_2$ compared to SOPP2 and SOPP3,[3] it was selected as the preferred LOV domain for LITag, and was termed LOV*. **B.** Apparatus design for blue light irradiation. We used two Kessil PR160L lamps (440 nm) at 100% power, which provide a high intensity of blue light (CAUTION: a proper eye protection is required). **C**. General workflow for LITag proximity labeling. Cells expressing the desired LOV* fusion protein are obtained by either transient or stable transfection. Cells are then washed twice and incubated with a solution of the probe in HEPES buffer (Live Cell Imaging Solution, ThermoFisher A14291DJ) for 30 min at 37 °C. Next, blue light irradiation is applied for a few seconds, followed by a downstream analysis of protein biotinylation. **D.** Testing different media for LITag. HEK293T cells expressing FLAG-LOV*-H2B were treated with BP probe (500 µM) in the indicated media and irradiated with blue light (3 s). Labeling efficiency analysis was performed as in panel A. The experiment indicates optically clear HEPES buffer to be ideal. The method is also compatible with phenol red-free DMEM culture media; the presence of phenol red considerably hampers the labeling. **E.** Testing various BP concentrations. Cells expressing FLAG-LOV*-H2B were treated with the indicated concentration of BP and then irradiated. Labeling was monitored by western blot as in panel A. Note: for this experiment, we employed longer irradiation times (minutes), as a different irradiation apparatus was used – a photoreactor that emits blue light with lower intensity compared to the Kessil PR160L lamps.[5] **F.** Cell viability assays (CellTiter Glo 2.0) with cells expressing MTS-LOV* (HEK293T) or PARP1-LOV* (U2OS). The PARP1-LOV*-expressing cells exhibited no photosensitivity even after 10 s irradiation. The MTS-LOV*-expressing cells showed some photosensitivity when irradiated for more extended periods (5 s or longer). Error bars – S.D., n = 3. Data were analyzed using a two-sample *t*-test, * $p < 0.05$, ns – not significant. **G.** Testing different biotin-containing probes for their LOV*-mediated activation.

HEK293T cells expressing FLAG-LOV*-H2B were treated with the indicated probe (500 µM). Labeling efficiency was monitored by western blot as a function of irradiation. BP and BA, which have a low redox potential (phenol: 0.633 V vs. SCE, aniline: 0.625 V vs. SCE)[6], undergo single electron transfer (SET) to the triplet-excited FMN (redox potential ~ 1.5 V vs. SCE[7]). On the other hand, aryl-azide and aryl-diazirine probes, which are activated by other photocatalysts through different mechanisms,[8,9] do not react with the triplet-excited FMN. Top – western blot showing protein biotinylation in cells expressing the LOV*-H2B fusion. Bottom – chemical structure of the various probes tested. **H.** Treatment of the cells with 1 mM $H_2O_2$, even when incubated for 1 min, enhances LITag efficiency. The effect was more pronounced for BP than for BA. HEK293T cells expressing FLAG-LOV*-H2B were incubated with the indicated probe (BP - 500 µM; BA, 250 µM) for 30 min. $H_2O_2$ (1 mM) was added to the cell medium at different time points during this incubation period. Labeling efficiency was monitored by western blot as a function of irradiation. **I.** $H_2O_2$ does not enhance LITag efficiency when performed *in vitro* with purified proteins. LOV* (10 µM), recombinantly expressed in *E. coli*, was mixed with BSA (1 mg/mL) in PBS. The solution was irradiated for the indicated time and biotinylation was analyzed by western blot. Ponceau staining of the membrane served as a loading control. **J.** Addition of glutathione to the *in vitro* reaction decreases the labeling efficiency. Reactions were performed as in panel I.

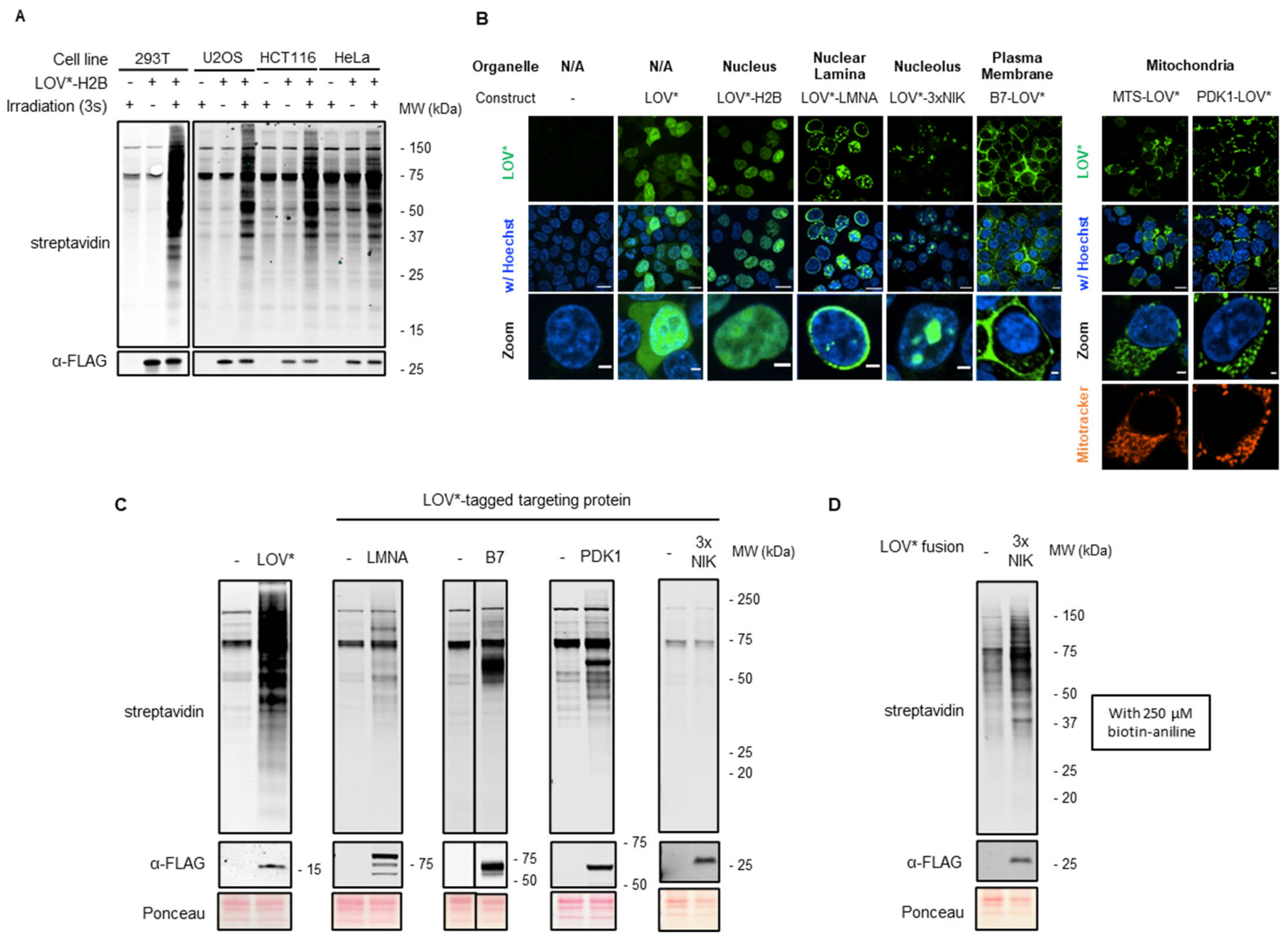


**Figure S2.** Scope of LITag. **A.** Testing of LITag in different cell lines. The indicated cells were transfected with a plasmid encoding for the FLAG-tagged LOV*-H2B fusion. After 24 h, the LITag workflow was employed, and biotinylation was analyzed by western blot as in Fig. S1A. The high transfection efficiency of HEK293T cells resulted in a considerably higher expression level of the LOV* fusion, thereby producing more protein tagging. **B.** Live-cell imaging showing LOV* targeting to different organelles in HEK293T cells. LOV* (green) was visualized by its intrinsic fluorescence. Mitochondria (orange) are labeled with MitoTracker Deep Red FM. Hoechst 33342 (blue) is a nuclear marker. Scale bar: second row, 10 μm; third row, 2 μm. **C.** Western blot showing protein biotinylation in different HEK293T organelles, in accordance with the immunofluorescence images in **Figure 1D**. Cells were treated with BP (500 μM) before blue light irradiation (3 s, except of LOV*-LMNA, where 1 s irradiation was used). Biotin signal could be detected under these conditions for all constructs except for the nucleolus targeted LOV* (3xNIK). Ponceau staining of the membrane served as a total protein loading control. **D.** Nucleolus protein labeling was achieved by switching to the biotin-aniline probe (250 μM).

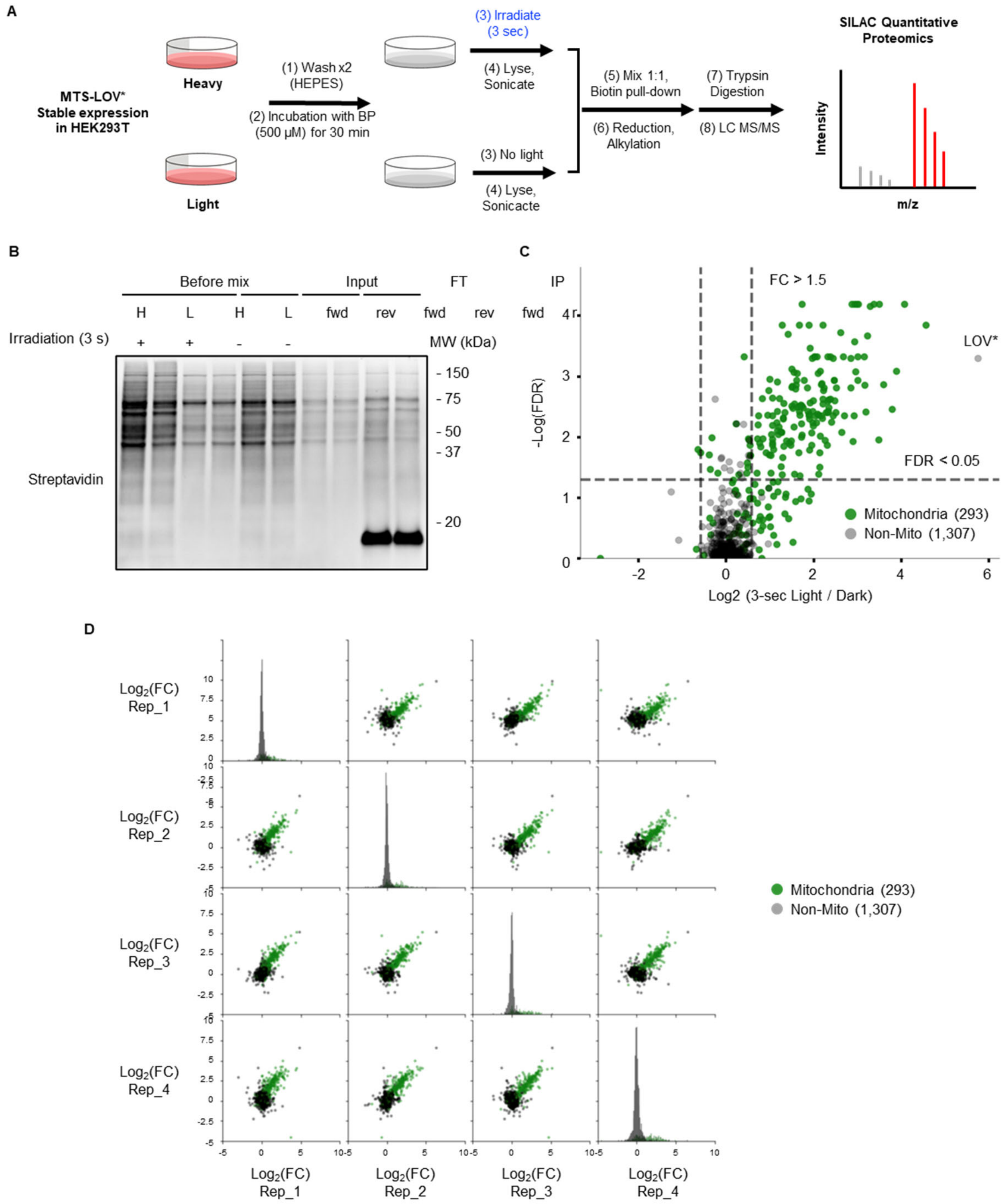


**Figure S3.** LITag labeling of the mitochondria. **A.** Workflow for SILAC-based proteomics. HEK293T cells stably expressing the MTS-LOV* fusion were cultured in "heavy" and "light" media. In the "forward" experiment, upon incubation with BP, live-cell LITag labeling was induced in the "heavy" cells by 3 s irradiation, while the "light" cells were kept in the dark. Subsequently, cells were lysed and sonicated to solubilize the entire proteome. Biotinylated proteins were pulled-down by streptavidin-coated magnetic beads. Following reduction and alkylation, proteins were digested on-beads and analyzed by LC-MS/MS. The experiment was repeated 4 times (2 "forward" and 2 "reverse"). **B.** Protein tagging in the samples used for SILAC-based proteomics was visualized by western blot. Two replicates were analyzed – one "forward" and one "reverse". **C.** Volcano plot displaying the SILAC-LITag data. Mitochondrial proteins are plotted in green (see Table S1 for the data). Thresholds are set at >1.5-fold change in enrichment and FDR <0.05. **D.** Replicates correlation plot. The $\log_2$ (FC) values obtained from each replicate were plotted against the $\log_2$ (FC) values of the other three replicates, showing overall good reproducibility of mitochondrial proteins enrichment.

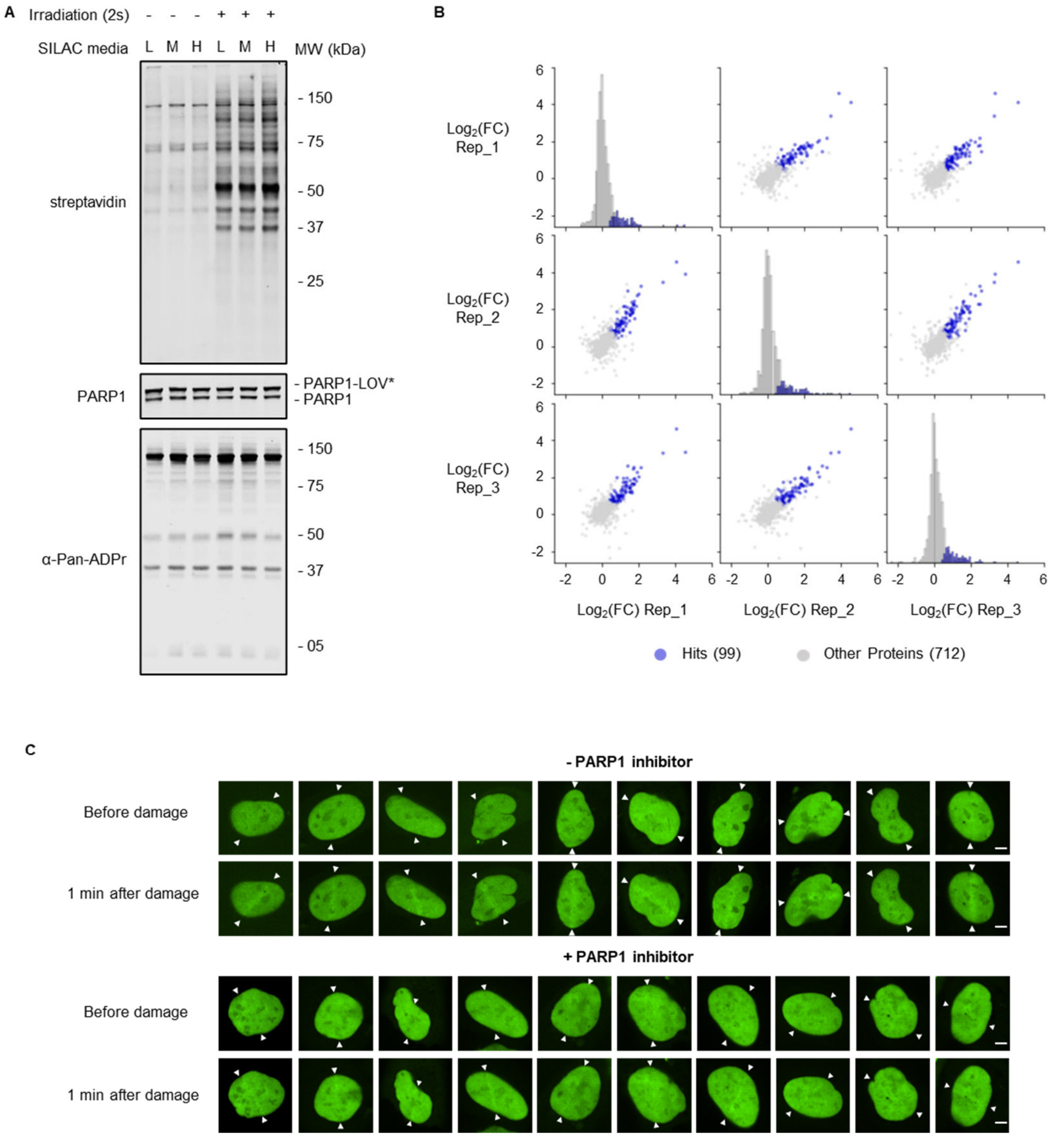


**Figure S4.** Investigation of PARP1 neighborhood following DNA damage. **A.** A stable U2OS cell line expressing a doxycycline-inducible PARP1-LOV* fusion was used in this experiment. Cells were incubated with doxycycline (400 ng/mL) for 12 h, upon which the expression level of PARP1-LOV* was similar to that of the endogenous PARP1. Cells were incubated for 30 min with BP (500 µM) in the presence of 1 mM $H_2O_2$ to induce DNA damage and protein poly(ADP-ribosylation) (PARylation). LITag was induced by 2 s irradiation, while the negative control was kept in the dark. This experiment employed "light" (L), "medium" (M) and "heavy" (H) SILAC media in three replicates according to the following settings: replicate 1 – 2 s irradiation (M) Vs. dark (L); replicate 2 – 2 s irradiation (H) Vs. dark (M); replicate 3 – 2 s irradiation (L) Vs. dark (H). The western blot shows protein biotinylation in the various samples used for proteomics using IRDye 800CW-labeled streptavidin. Western blot against PARP1 was used as a loading control, while an anti-pan-ADPr probe was used to show activation of PARP1 PARylation activity. **B.** Replicates correlation plot. The $\log_2$ (FC) values obtained from each replicate were plotted against the $\log_2$ (FC) values of the other two replicates. **C.** Laser microirradiation and live-cell imaging of eGFP-tagged HMGB1 in U2OS cells; supplementary images for Figure 3F. Cells were incubated for 1 h with or without PARP1 inhibitor (talazoparib, 250 nM), followed by laser-induced DNA damage. HMGB1 accumulation at the damage site is shown 1 min after laser microirradiation. Scale bar, 5 µm.

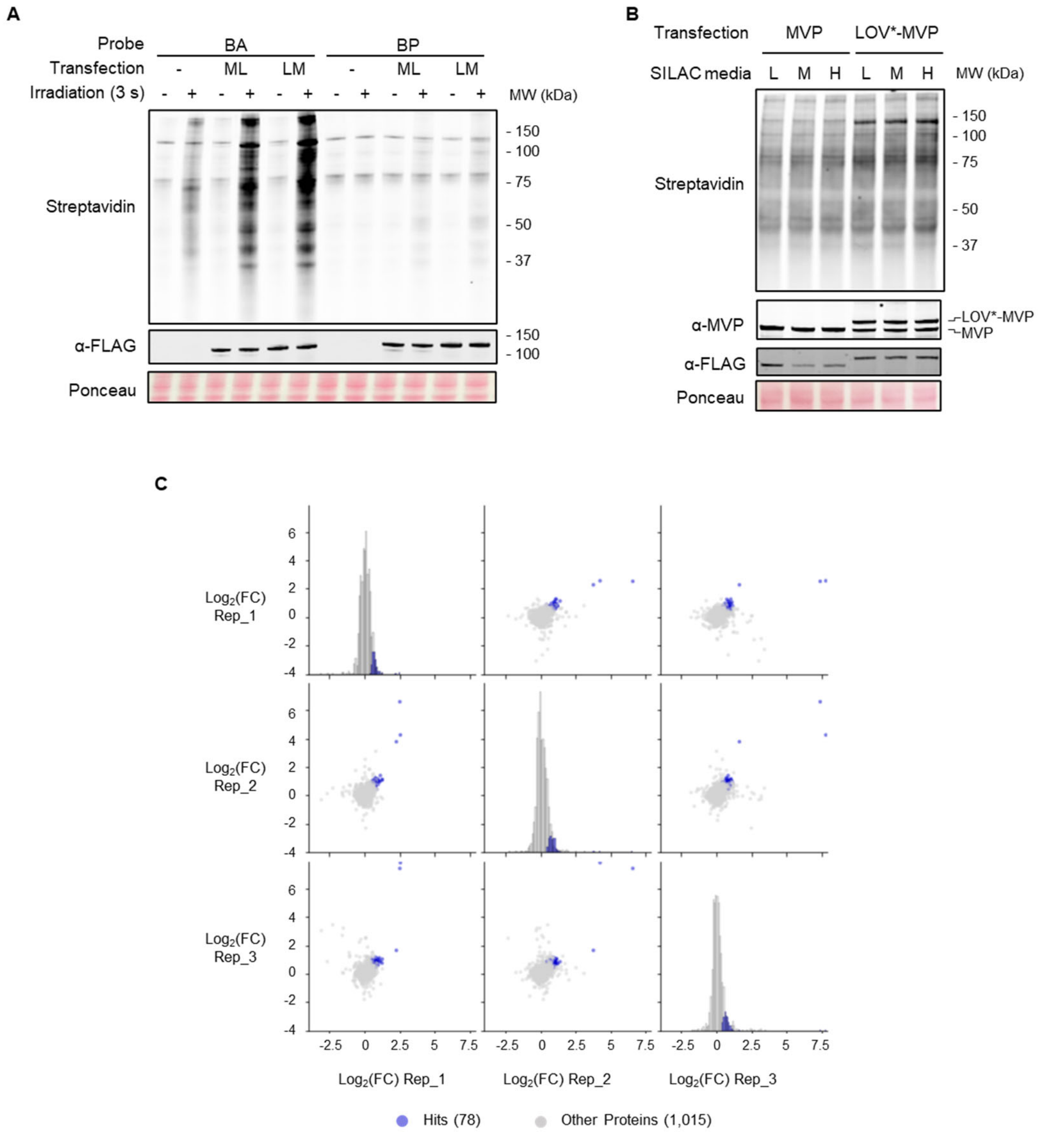


**Figure S5.** Studying the interactome of the Major Vault Protein (MVP). **A.** Western blot analysis showing protein biotinylation in HeLa cells following transient transfection with LOV*-tagged MVP, probe incubation and light irradiation. Incubation with BA (250 µM) afforded robust protein tagging, while the use of the BP probe (500 µM) afforded weaker labeling in this case. ML, MVP-LOV*; LM, LOV*-MVP. Labeling efficiency was monitored by western blot as a function of irradiation. Ponceau staining of the membrane served as a total protein loading control. **B.** For SILAC-based proteomics, HeLa cells, grown in SILAC media, were transfected with FLAG-tagged LOV*-MVP (negative control – transfection with FLAG-tagged MVP). 20 h post transfection, cells were incubated for 30 min with BA (250 µM) and irradiated for 3 s to induce protein tagging. This experiment employed "light" (L), "medium" (M) and "heavy" (H) SILAC media in three replicates according to the following settings: replicate 1 – LOV*-MVP (M) Vs. MVP (L); replicate 2 – LOV*-MVP (H) Vs. MVP (M); replicate 3 – LOV*-MVP (L) Vs. MVP (H). Western blot is showing protein biotinylation in the various samples used for proteomics. **C.** Replicates correlation plot. The $\log_2$ (FC) values obtained from each replicate were plotted against the $\log_2$ (FC) values of the other two replicates.

**Table S1.**

**Proteomics data of the LITag mitochondria labeling.** The table includes a list of the enriched proteins (FC > 1.5, FDR<0.05) and a comprehensive list of all the proteins in the datasets, with their FC and FDR. Mitochondrial and submitochondrial localization of the detected proteins is also detailed.

**Table S2.**

**Proteomics data of the LITag PARP1 interactome labeling.** The table includes a list of the enriched proteins (FC > 1.5, FDR<0.05) and a comprehensive list of all the proteins in the datasets, with their FC and FDR. Information regarding the PARylation, PAR binding, and RNA binding of the enriched proteins is also detailed.

**Table S3.**

**Proteomics data of the LITag MVP interactome labeling.** The table includes a list of the enriched proteins (FC > 1.5, FDR<0.05) and a comprehensive list of all the proteins in the datasets, with their FC and FDR. The enriched proteins were categorized manually based on their cellular function.

**Movie S1.**

**The PARP1-LOV* fusion is recruited rapidly to the site for DNA damage.** Live-cell imaging of the PARP1-LOV* green fluorescence, tracking the recruitment of the fusion to the line of DNA damage in real-time.

**Material and Methods**

Antibodies:

| Antibody (Host) | Usage and Dilution | Source | Catalog Number |
|---|---|---|---|
| anti-FLAG (mouse) | Western blot – 1:1500<br>Immunofluorescence – 1:500 | Sigma | F1804 |
| anti-PARP1 (rabbit) | Western blot – 1:1500 | Abcam | ab227244 |
| anti-Pan-ADP-ribose (rabbit) | Western blot – 1:1500 | Sigma | MABE1016 |
| anti-MVP (rabbit) | Western blot – 1:1500 | ThermoFisher | 16478-1-AP |
| anti-PARP4 (rabbit) | Western blot – 1:1500 | Abcam | ab133745 |
| anti-BAG2 (rabbit) | Western blot – 1:1500 | ThermoFisher | PA5-30922 |
| anti-GAPDH (mouse) | Western blot – 1:1500 | ThermoFisher | MA1-16757 |
| IRDye® 680RD –<br>anti-Mouse IgG (goat) | Western blot – 1:10000 | LI-COR | 926-68070 |
| IRDye® 800CW –<br>anti-Rabbit IgG (goat) | Western blot – 1:10000 | LI-COR | 926-32211 |
| IRDye® 800CW Streptavidin | Western blot – 1:10000 | LI-COR | 926-32230 |
| anti-Mouse IgG (goat) –<br>Alexa Fluor™ 488 | Immunofluorescence – 1:1000 | ThermoFisher | A-11001 |
| NeutrAvidin™, Rhodamine<br>Red™-X conjugate | Immunofluorescence – 1:500 | ThermoFisher | A6378 |

**Cloning**

DNA constructs:

| Name | Fusion components | DNA sources and references |
|---|---|---|
| LOV*-H2B in pCMV | FLAG-LOV*-H2B | LOV* – gBlock®; H2B – Muir lab plasmids[10] |
| miniSOG-H2B in pCMV | FLAG-miniSOG-H2B | miniSOG – L102Q point-mutation of LOV* |
| miniSOG2-H2B in pCMV | FLAG-miniSOG2-H2B | miniSOG2 – gBlock® |
| SOPP2-H2B in pCMV | FLAG-SOPP2-H2B | SOPP2 – W81L and L102V point mutations of LOV* |
| SOPP3-H2B in pCMV | FLAG-SOPP3-H2B | SOPP3 – gBlock® |
| LOV* in pCMV | FLAG-LOV* | |
| LOV*-LMNA in pCMV | FLAG-LOV*-LMNA | LMNA – cloned from 'V5-APEX2-LMNA' (Addgene plasmid # 129275), a gift from Alice Ting |
| PDK1-LOV* in pCMV | PDK1-FLAG-LOV* | PDK1 – cloned from 'pWZL Neo Myr Flag PDK1' (Addgene plasmid # 20564), a gift from William Hahn & Jean Zhao |
| LOV*-3xNIK in pCMV | FLAG-LOV*-3xNIK | 3xNIK – nucleolus localization signal from NF-κB inducing kinase (NIK)[11] – insertion with PCR primers |
| LOV*-B7 in pCMV | Igκ-FLAG-LOV*-$B7_{143\text{-}306}$ | Igκ leader sequence – insertion with PCR primers; Mouse B7 (CD80 antigen), residues 143-306[12] – gBlock® |
| MTS-LOV* in pLenti-EF1a | MTS-FLAG-LOV* | mitochondrial targeting sequence (MTS) from COX4 – insertion with PCR primers |
| PARP1-LOV* in PB-TRE3G | PARP1-FLAG-LOV* | PARP1 – Muir lab plasmids[13] |
| LOV*-MVP in pCMV | FLAG-LOV*-MVP | MVP – cloned from 'pDONR223_MVP_WT_V5' (Addgene plasmid # 82977), a gift from Jesse Boehm & Matthew Meyerson & David Root |
| MVP-LOV* in pCMV | MVP-FLAG-LOV* | |
| LOV* in pET30 | 6xHis-LOV* | |
| eGFP-HMGB1 in pcDNA3.1 | FLAG-eGFP-HMGB1 | HMGB1 – eGFP was cloned into the 'pcDNA3.1 Flag hHMGB1' vector (Addgene plasmid 31609), a gift from Yasuhiko Kawakami |

All constructs used in this study were generated using standard molecular cloning procedures (Gibson assembly, restriction cloning, PCR coupled with blunt-end ligation) and confirmed by Sanger sequencing. Plasmid maps and protein sequences for all constructs used in this study are given below (pages S20-S35).

**General Laboratory Methods**

Common reagents and chemicals were purchased from MilliporeSigma unless stated otherwise and were used without further purification. Oligonucleotide primers for cloning were purchased from Integrated DNA Technologies (IDT) or MilliporeSigma. gBlock® Gene Fragments were synthesized by IDT. PrimeSTAR® HS DNA Polymerase (premix, Takara R040A) was used for PCR. Gibson Assembly Master Mix was purchased from New England Biolabs (E2611L). BL21(DE3) chemically competent *E. coli* cells and Subcloning Efficiency DH5α competent *E. coli* cells were generated in-house from cells purchased from Invitrogen. DNA purification kits for plasmid purification were purchased from QIAGEN. All plasmid sequences were verified by Sanger sequencing performed by GENEWIZ. Acrylamide, TEMED, APS, and Criterion Empty Cassette were purchased from Bio-Rad. Nitrocellulose membrane (0.45 µm) for western blot was purchased from ThermoFisher (88018). Processing of fluorescence microscopy images was performed using ImageJ2 (National Institutes of Health). Statistical analyses were conducted in Prism v.9.2.0.

**Cell culture**

HEK293T, U2OS, HeLa, A549, and HCT116 were cultured in Dulbecco's Modified Eagle Medium (DMEM, Thermo Fisher 11995-005), supplemented with 10% v/v FBS (Atlanta Biologicals S12450H), 100 U/mL penicillin, 100 µg/mL streptomycin (ThermoFisher 15140-122), and 2 mM L-Glutamine (Thermo Fisher 25030-081). Cells were maintained in an incubator at 37 °C with 5% $CO_2$. For HEK293T, plates were pre-coated with 0.01 % poly-l-lysine solution (Sigma P8920).

**Transfection and construction of stable cell lines**

Transfection for transient protein expression in mammalian cells was performed using Lipofectamine™ 2000 Transfection Reagent (Thermo Fisher 11668-019) following the manufacturer's instructions. For generating the MTS-LOV* HEK293T stable cell line, we used lentivirus transduction. 48 h post-transduction, cells were cultured with puromycin (ThermoFisher A1113803, 1 µg/mL) for 5 days to select for transduced cells. For generating the PARP1-LOV* U2OS stable cell line, cells were transfected with a Piggybac transposon plasmid encoding for PARP1-LOV* under the control of a doxycycline-inducible promoter, and a Piggybac transposase plasmid (a gift from the Kadoch lab, Dana Farber Cancer Institute) (2:1 ratio). Lipofectamine™ 3000 Transfection Reagent (ThermoFisher L3000015) was used for this transfection according to the manufacturer's instructions. 30 h after transfection, cells were cultured in the presence of 10 µg/mL blasticidin S-HCl (BCD, Thermo Fisher A11139-03) for 7 days. The BCD-selected cells were further sorted based on positive green fluorescence following 48 h incubation with doxycycline (400 ng/mL) (Stem Cell Technologies 72742, 2 mg/mL stock solution in DMSO) using the BD FACSAria Fusion instrument. LITag and live-cell imaging experiments in this paper were carried out after incubation with doxycycline (400 ng/mL) for 12 hours.

**Live cell imaging**

For visualizing the LOV* construct in live cells, cells were stained with Hoechst 33342 dye (Invitrogen H3570, 1:2000 dilution) for 5 min and subsequently washed twice with Live Cell Imaging Solution (ThermoFisher A14291DJ). To show the mitochondrial localization of PDK1-LOV* and MTS-LOV*, cells were also stained with MitoTracker Deep Red FM

(ThermoFisher M22426, 100 nM) for 30 min at 37 °C under 5% $CO_2$. Cells were imaged using a Nikon CSU-21 spinning disk confocal microscope with NIS Elements (version 5.2) software. We used a 488 nm laser for the excitation of LOV*, 405 nm for the excitation of Hoechst 33342, and 561 nm for Mitotracker excitation. Images were taken with a 60x Plan Apo λ Oil objective and Hamamatsu ORCA-Flash4.0 sCMOS camera.

**Laser Microirradiation**

For the PARP1-LOV* laser microirradiation experiment, the PARP1-LOV* U2OS stable cell line was incubated with doxycycline (400 ng/mL) for 12 h. Next, cells were sensitized by incubation with Hoechst 33342 dye (1:2000 dilution) for 5 min and subsequently washed twice with Live Cell Imaging Solution (ThermoFisher, A14291DJ). Nuclei irradiation was performed using an OBIS LX 405 nm 100 mW Laser (30% laser power, 2000 µs dwell time) connected to a Nikon Galvo Miniscanner. Images were acquired using a confocal microscope with the above specifications.

For the eGFP-HMGB1 laser microirradiation experiment, U2OS cells were transfected with the eGFP-HMGB1 construct. After 24 h, cells were sensitized, irradiated, and imaged as described for the PARP1-LOV* cells. When PARP1 inhibitor was included, cells were incubated with talazoparib (MedChemExpress HY-16106, 250 nM, diluted from a 5 mM stock solution in DMSO) for 1 h in culture media before microirradiation. Talazoparib was kept in solution while staining with Hoechst 33342 and in the final imaging solution. For quantifying the recruitment of eGFP-HMGB1 to the DNA damage site, we defined a Region of Interest (ROI) in ImageJ2 along the stimulation line (width – 20 pixels, resolution – 13.3 pixels/µm). The relative fluorescence intensity of the ROI was calculated by dividing the intensity of this ROI at the indicated time points after DNA damage by the intensity of the same ROI before DNA damage. Data are presented as the average of 21 irradiated cells, ± SD.

**Cell viability assay**

To probe the photosensitivity of cells expressing LOV* fusions, we used the CellTiter-Glo® 2.0 Cell Viability Assay (Promega G9241). Cells were seeded in a 24-well plate with black walls and clear bottom ($5x10^4$ cells/well); for the PARP1-LOV* U2OS cell line, incubation was performed in the presence of doxycycline (400 ng / mL). After 24 h, cells were irradiated with blue light for the indicated periods (1-10 s) and further incubated for 24 h. Then, the culture media was aspirated and replaced with fresh DMEM at RT (200 µL / well), to which the CellTiter-Glo® 2.0 solution (200 µL/well) was added. Following orbital shaking for 2 min and incubation for an additional 10 min at RT, luminescence was measured using the SpectraMax M3 Microplate Reader. Data are presented as the average of three replicates ± SD.

**LITag labeling**

To perform LITag experiments, cells were washed twice with PBS (137 mM NaCl, 12 mM Phosphate, 2.7 mM KCl, pH 7.4) or Live Cell Imaging Solution. A solution of biotin-phenol (BP, ApexBio A8011, 500 µM, 500x dilution from a 250 mM solution in DMSO) or biotin-aniline (BA, Iris Biotech LS-3970, 250 µM, 1000x dilution from a 250 mM solution in DMSO) in Live Cell Imaging Solution was added, and the cells were incubated for 30 mins at 37 °C under 5% $CO_2$. For the data shown in Figure S1G, we also tested N-[1-(4-azido-2,3,5,6-tetrafluorophenyl)-1-oxo-6,9,12-trioxa-2-azapentadec-15-yl]-

biotinamide (TFPA-PEG$_3$-Biotin, 500 μM, ThermoFisher 21303) and N-(3-oxo-1-(4-(3-(trifluoromethyl)-3H-diazirin-3-yl)phenyl)-6,9,12-trioxa-2-azatetradecan-14-yl)-biotinamide (biotin-PEG$_3$-TDBnA, 500 μM, ref [9]). Following probe incubation, cells were irradiated using two Kessil PR160L lamps (440 nm, see Figure S1B) for the indicated time. Subsequently, cells were washed twice with PBS, and further processing was performed according to the various methods described below.

**Western Blot**

Following LITag labeling, 1xSDS sample loading buffer (100 mM Tris-Cl, pH 6.8, 3% SDS, 15% Glycerol, 2.25% β-mercaptoethanol, 0.015% bromophenol blue) was added to lyse the cells. Samples were boiled at 98°C for 30 mins and were subsequently loaded on a 10% bis-tris polyacrylamide gel. Proteins were transferred to a nitrocellulose membrane which was then blocked with 5% w/v nonfat dry milk in TBST (20 mM Tris-Cl, pH 7.6, 150 mM NaCl, 0.1% v/v Tween-20) for 30 mins at room temperature. Next, membranes were incubated with the indicated primary antibodies (diluted in TBST containing 1% BSA) at room temperature for 2 h or at 4 °C overnight. After washing three times with TBST, the appropriate Li-Cor IRDye secondary antibodies (1:10000 dilution in TBST) were applied for 1 h at room temperature. For the detection of the biotin signal produced by LITag, membranes were incubated with an IRDye® 800CW-labeled streptavidin (LI-COR 926-32230, 1:10000 dilution in TBST) for 1 h at room temperature. Following incubation with the dye-labeled secondary antibodies, membranes were washed 3 times with TBST and imaged with Li-Cor Odyssey Infrared Imaging System.

**Immunofluorescence**

Cells were cultured in either a 35 mm glass-bottom petri dish (MatTek P35G-1.5-14-C) or a 24-well glass-bottom plate (Cellvis P24-1.5H-N). LITag labeling was performed at 30-40% confluency as described above. Subsequently, cells were fixed with 4% formaldehyde for 10 min, then permeabilized with 0.5% Triton in PBS for 15 min at room temperature. 3% BSA or 2.5% goat serum in PBS were used for blocking for at least 1 hour at room temperature. For experiments that only need biotin staining, cells were incubated with NeutrAvidin™-Rhodamine Red™-X (2 μg/mL, ThermoFisher A6378) and Hoechst 33342 (10 μg/mL) for 1 h at RT, washed 3 times with PBS, and imaged using a confocal microscope, with specifications as described above for live cell imaging. When staining with a primary antibody was also needed, the antibody was incubated with the cells overnight at 4 °C, followed by staining with a secondary antibody, NeutrAvidin™-Rhodamine Red™-X, and Hoechst 33342. Cells were washed 3 times with PBS and imaged using a confocal microscope with the above specifications.

**SILAC-based proteomics**

<u>General protocol</u> – "heavy", "medium", and "light" media were prepared using 'DMEM for SILAC' (ThermoFisher 88364) supplemented with 10% dialyzed FBS (ThermoFisher A3382001), 100 U/mL penicillin, 100 μg/mL streptomycin (ThermoFisher 15140-122), and the appropriate amino acids: "heavy" (H) media – L-lysine-$^{13}C_6$,$^{15}N_2$ dihydrochloride (Cambridge Isotope Laboratories (CIL) CNLM-291-H) and L-arginine-$^{13}C_6$,$^{15}N_4$ hydrochloride (CIL CNLM-539-H); "medium"

(M) media – L-lysine-4,4,5,5-$d_4$ dihydrochloride (CIL DLM-2640) and L-arginine-$^{13}C_6$ hydrochloride (CIL CLM-2265-H); "light" (L) media – L-lysine hydrochloride (Sigma L1262) and L-arginine hydrochloride (Sigma A3909). Cells were cultured in SILAC media for at least 8 doublings to ensure complete isotopic labeling. For the LITag labeling step, cells were seeded in 6-well plates; labeling was performed at 95-100% confluency. We used a total of $2x10^7$ cells per experimental replicate ($1x10^7$ cells for each SILAC condition). After LITag labeling (as described above), the cells were washed twice with PBS, followed by an additional wash in PBS for 5 min. The cells were incubated with lysis buffer (10 mM tris, pH 8.0, 100 mM NaCl, 1 mM EDTA, 0.5 mM EGTA, 0.5% sodium lauryl sarcosine, 0.1% sodium deoxycholate, 1x Halt protease inhibitor cocktail, ThermoFisher 78438; 150 µL/well) for 10 min at RT and were subsequently transferred to a 5 mL Eppendorf tube. The cell lysate was sonicated on ice using Fisherbrand 505 Sonic Dismembrator (3 mm microtip, 25% amplitude, 4x15 s), and the solution was clarified by centrifugation (10 min at 4 °C, 17000 g). The protein concentration was determined by BCA (1-2 mg/mL), and samples were diluted with the appropriate volume of lysis buffer so that the concentration of the different L/M/H samples was even. SILAC samples were mixed 1:1 and diluted with 1 volume of binding buffer (0.25% NP-40 alternative in TBS, pH 7.6). Streptavidin magnetic beads (Streptavidin Mag Sepharose, Cytiva 28985799, 300 µL slurry), pre-equilibrated with binding buffer, were added, and the samples were rotated head-over-head for 2 h at RT. Subsequently, the beads were collected and washed with binding buffer (once), 1% SDS in PBS (twice), 1 M NaCl in PBS (twice), and 100 mM ammonium bicarbonate (three times). The beads were suspended in 300 µL of 5 mM DTT in 100 mM ammonium bicarbonate and mixed at 56 °C for 30 min. The suspension was cooled down to RT, and 9 µL of 500 mM iodoacetamide in water was added, followed by rotation at RT for 20 min in the dark. The beads were washed 3 times with 50 mM ammonium bicarbonate and then resuspended in 300 µL 50 mM ammonium bicarbonate containing 3 µg of trypsin gold (Promega V5280). Following rotation at 37 °C overnight, the supernatant was collected and dried using a SpeedVac vacuum concentrator. The samples were then subjected to LC-MS/MS analysis (see below).

Specific experimental details:

1. MTS-LOV* (n=4): 2 forward experiments – 3 s irradiation (H) vs. no irradiation (L); 2 reverse experiments – 3 s irradiation (L) vs. no irradiation (H).

2. PARP1-LOV* (n=3): replicate 1 – 2 s irradiation (M) vs. dark (L); replicate 2 – 2 s irradiation (H) vs. dark (M); replicate 3 – 2 s irradiation (L) vs. dark (H). Doxycycline (400 ng/mL) was added 12 h prior to labeling. 3. LOV*-MVP (n=3): replicate 1 – LOV*-MVP (M) vs. MVP (L); replicate 2 – LOV*-MVP (H) vs. MVP (M); replicate 3 – LOV*-MVP (L) vs. MVP (H). 20 h before labeling, cells were transfected with the appropriate expression plasmid (LOV*-MVP or MVP), using Lipofectamine™ 2000 Transfection Reagent (Thermo Fisher 11668-019); 1.5 µg of DNA with 4.5 µL Lipofectamine 2000 per 6-well plate.

**Mass Spectrometry**

Following desalting and drying the trypsin-digested samples, the peptides were resuspended in 21 µL of 0.1% formic acid (pH 3.0), and 2 µL was injected per run using an Easy-nLC 1200 UPLC system. The peptides were separated on a 45 cm × 100 µm capillary column packed with 1.9 µm C18-AQ, mated to a metal emitter in-line with an Orbitrap Fusion Lumos (Thermo Scientific). The column temperature was set to 50 °C, and a 2 h gradient method with a flow rate of 360 nL/min was used. Mass spectrometers were operated in data-dependent and positive ionization mode. MS1 spectra were recorded at a resolution of 120k using an automatic gain control (AGC) target value of $4x10^5$, maximum injection time (maxIT) of 50 ms, and a range of 375-1500 *m/z*. After peptide fragmentation via HCD (35% collision energy), MS2 spectra were acquired using an AGC target value of $1 \times 10^4$ and a maxIT of 54 ms.

**Data Processing**

Data processing was performed using MaxQuant (v2.1.3.0)[14] with default parameters unless noted otherwise. Match between runs was selected. The Uniprot human proteome reference database was used (proteome ID UP000005640) as a reference. Trypsin/P was selected as the enzyme. Carbamidomethylation of cysteine was specified as a fixed modification. Methionine oxidation and N-terminal acetylation were set as variable modifications. From the "ProteinGroups" file generated by MaxQuant, proteins identified as common contaminants, identified only by sites, and proteins matched in the reverse decoy database were removed. Only proteins identified in all experimental replicates were reported.

Imputation: Some proteins were identified in all replicates of one experimental condition but not in all the replicates of the other condition (e.g., proteins were identified in all replicates in the '3 s irradiation' condition but were not detected in all replicates in the 'dark' condition). For those proteins, we employed imputation according to the following method. Peptide intensities of these proteins were extracted from the "Peptides" file generated by MaxQuant. The missing value was substituted with the $10^{th}$ percentile of the intensities detected in the specific SILAC condition, and the SILAC ratio was calculated for all the peptides. A protein's SILAC ratio was taken from the median ratio of all assigned unique peptides. A python script was written for the purpose of imputation and ratio calculation.

Normalization: For the PARP1-LOV* LOV*-MVP LITag experiments, the protein SILAC ratios were normalized based on the median of the distribution of ratios for a specific SILAC channel (e.g., H/L); the ratio for each detected protein was divided by this median to normalize. In the case of the MTS-LOV* experiment, only the non-mitochondrial proteins were used to generate the median,[15] as these proteins are not expected to be labeled and therefore should produce similar MS signal with and without irradiation.

Statistical Analysis: Following the transformation of the SILAC ratios to the $\log_2$ scale, a one-sample *t*-test was employed with the null hypothesis that $Log_2(FC) = 0$. The *t*-test p-values were adjusted using the Benjamini–Hochberg procedure to obtain the FDR values. A protein with an FC>1.5 and an FDR<0.05 was considered significant.

**Mitochondrial mRNA enrichment**

For mitochondrial RNA labeling, MTS-LOV*-expressing HEK293T cells were cultured in two 6-well plates (4 wells per plate). We performed the LITag protocol described above, where the 4 wells in one plate were irradiated for 3 s, and the other 4 wells were kept in the dark. Following washing with PBS, RNA was extracted separately from each well using Qiagen's RNeasy Mini kit (Qiagen 74104), providing 4 replicates per experimental condition. About 30 µg of RNA was obtained from each well, and 15 µg was used for further processing. Biotinylated RNA molecules were pulled down according to a previously published protocol.[16] We used Pierce streptavidin magnetic beads (ThermoFisher 88816), 10 µL beads for 15 µg of RNA. The beads were washed 3 times in wash buffer (5 mM Tris-HCl, pH 7.5, 0.5 mM EDTA, 1 M NaCl, 0.1% Tween-

20), followed by one wash in 0.1 M NaCl solution. The washed beads were then suspended in 100 µL 0.1 M NaCl and incubated with 100 µL RNA (diluted in water) on a rotator for 2 h at 4°C. The beads were placed on a magnet, and the supernatant was discarded. The beads were washed 3 times in wash buffer and resuspended in 54 µL water. A 3X proteinase digestion buffer was made: 1.1 mL buffer containing 330 µL 10X PBS pH = 7.4 (ThermoFisher AM9624), 330 µL 20% N-Laurylsarcosine sodium salt solution (Sigma L7414), 66 µL 0.5 M EDTA, 16.5 µL 1 M DTT and 357.5 µL water. 33 µL of this 3X proteinase buffer was added to the beads along with 10 µL Proteinase K (20 mg/mL, Thermofisher AM2546) and 3 µL Ribolock RNase inhibitor (ThermoFisher EO0381). The beads were transferred to PCR tubes and incubated in a thermocycler at 42°C for 1 h, followed by 55°C for 1 h. The supernatant was collected, and the RNA was purified using the RNA Clean & Concentrator-5 kit (Zymo Research R1013). Elution was done in 10 µL.

The eluted RNA was converted to cDNA using the High-Capacity cDNA Reverse Transcription Kit (ThermoFisher 4368814) according to the manufacturer's instructions. In addition, 1 µg of RNA before biotin pull-down (input) was converted to cDNA. The resulting cDNA samples were tested in an RT-qPCR experiment using the SYBR™ Green PCR Master Mix (ThermoFisher 4309155) in a ViiA 7 Real-Time PCR System. The following primers were used:

MT-CO2: forward – AACCAAACCACTTTCACCGC; reverse – CGATGGGCATGAAACTGTGG.

MT-ND1: forward – CACCTCTAGCCTAGCCGTTT; reverse – CCGATCAGGGCGTAGTTTGA.

GAPDH: forward – TGTCAAGCTCATTTCCTGGTAT; reverse – CTCTCTTCCTCTTGTGCTCTTG.

XIST: forward – CCCTACTAGCTCCTCGGACA; reverse – ACACATGCAGCGTGGTATCT.

The dCt for each condition (3 s irradiation or dark) was calculated by subtracting the input Ct from the pull-down Ct. LITag enrichment was assessed by the ddCt values [dCt (3 s irradiation) - dCt (dark)]. To determine the statistical significance of enrichment, a one-sample *t*-test was employed with the null hypothesis that ddCt = 0.

**Expression of recombinant LOV***

For the *in vitro* experiments described in Figure S1I and Figure S1J, we used a recombinant LOV* protein expressed in *E. coli*. BL21(DE3) cells were transformed with a plasmid encoding for the LOV* domain, containing a 6xHis tag at the N terminus. The cells were plated on a kanamycin plate (50 µg/mL). A colony was picked, and the cells were grown in a 5 mL LB broth medium with kanamycin (50 µg/mL) until turbidity was observed. Cells were transferred to 2x1 L flasks and incubated in LB broth medium (w/ kanamycin) at 37 °C until OD was 0.6. Flasks were cooled to 18 °C, and protein expression was induced with 0.5 mM IPTG. After overnight incubation at 18 °C, cells were collected by centrifugation at 4000 g for 20 min, resuspended in 30 mL of lysis buffer (20 mM Tris, pH 8, 500 mM NaCl, 1 mM PMSF), and lysed by sonication on ice using Fisherbrand 505 Sonic Dismembrator (3 mm microtip, 30% amplitude, 8x20 s). The cell lysate was cleared by centrifugation at 4 °C at 39000 g for 15 min. The supernatant was incubated for 1 h at 4 °C with 2 mL of HisPur™ Ni-NTA Resin (ThermoFisher 88221), pre-equilibrated with lysis buffer. The beads were then washed with 50 mL of lysis buffer supplemented with 20 mM imidazole, and the protein was eluted with 250 mM imidazole in lysis buffer (10 mL). The solution was concentrated to 1.5 mL by ultrafiltration using Amicon® Ultra-15 (Millipore UFC9010). Purification by gel filtration was performed on a HiLoad® 16/600 S75 column, pre-equilibrated with purification buffer (20 mM Tris, pH 8, 150 mM NaCl, 10% glycerol). The gel filtration fractions were analyzed by SDS-PAGE, and the major fractions containing

LOV* were pooled, concentrated down to 150 µM, aliquoted, and stored at -80 °C. LOV* has maximum absorption at 440 nm, with an extinction coefficient of 14500 $M^{-1}cm^{-1}$,[2] which was used to determine the protein concentration. The purified protein was analyzed by electrospray ionization quadrupole time-of-flight (ESI-Q-TOF) mass spectrometry using Bruker's micrOTOF-Q II. Calculated molecular weight: 13568.3 Da, observed: 13567.7 Da. See page S36 for LOV* mass spectrum and SDS-PAGE.

**In vitro experiments with recombinant LOV***

LITag labeling *in vitro* was performed using 10 µM of recombinant LOV* mixed with BSA (1 mg/mL) in PBS, pH 7.4, in the presence of 500 µM BP. The experiment was performed in a transparent 96-well plate with a 100 µL reaction volume. 50 µL of a 2x LOV* solution (20 µM) in PBS was placed in the reaction well, and 50 µL of a 2x BSA-BP solution (BSA, 2 mg/mL; BP, 1000 µM) in PBS was added. The mixture was incubated in the dark for 30 min at room temperature before blue light irradiation. For testing the effect of $H_2O_2$ on labeling efficiency, 1 µL of a 100x solution of $H_2O_2$ in water was added at various time points (30 min or 1 min before irradiation). For testing the effect of glutathione, the aforementioned 2x BSA-BP solution also contained the appropriate amount of 2x glutathione. Following irradiation with blue light, the samples were diluted with a 4xSDS loading dye and analyzed by western blot as described above.

**HeLa cell differential centrifugation**

To determine whether the LOV*-tagged MVP fusions incorporate in the vault particles, a differential centrifugation experiment was employed. HeLa cells in 60 mm dishes (95% confluency) were transfected with the various MVP plasmids. 20 h after transfection, cells were washed with cold PBS, harvested by scraping in 1 mL of cold PBS, centrifuged at 4 °C at 200 g for 5 min, and then resuspended in 200 µL of cold lysis buffer (50 mM tris, pH 7.4, 75 mM NaCl, 1.5 mM $MgCl_2$, 0.5% NP-40 alternative). Cells were incubated on ice for 10 min and then centrifuged at 17,000 g at 4 °C for 10 min. 30 µL of the cell lysate was kept as a reference for western blot, and the rest of the supernatant was transferred to a 5 mL open-top tube (Beckman Coulter 344057). Centrifugation was performed at 127000 g for 2 h at 4 °C, using the Beckman Optima™ L-80 XP ultracentrifuge with a Ti-55 rotor. The supernatant was transferred to a fresh 1.5 mL tube, and the pellet was resuspended in 1x SDS loading dye (170 µL). Samples were boiled and analyzed by western blot as described above.

**Sucrose gradient centrifugation**

For the sucrose gradient centrifugation experiment, we used a confluent 100 mm dish of HeLa or A549 cells. Cells were washed with cold PBS, harvested by scraping in 1.5 mL of cold PBS, and centrifuged at 4 °C at 200 g for 5 min. Cells were resuspended in 250 µL of cold buffer A (50 mM Tris-Cl, pH 7.4, 1.5 mM $MgCl_2$, 75 mM NaCl) containing 1% NP-40 alternative, 15% glycerol, 10% sucrose, and 1x Halt protease inhibitor cocktail, incubated on ice for 10 min, and centrifuged at 17,000 g for 10 min. The soluble lysate was applied to a 30-60% sucrose step gradient (made in Buffer A in a 5 mL open-top tube, Beckman Coulter 344057) and centrifuged at 90,000 g overnight at 4 °C, using the Beckman Optima™ L-80 XP ultracentrifuge with a Ti-55 rotor. Gradient fractions (250 µL each) were collected, diluted with 4x SDS loading dye, boiled, and analyzed by western blot as described above.

**DNA Constructs**

<u>LOV*-H2B in pCMV</u>

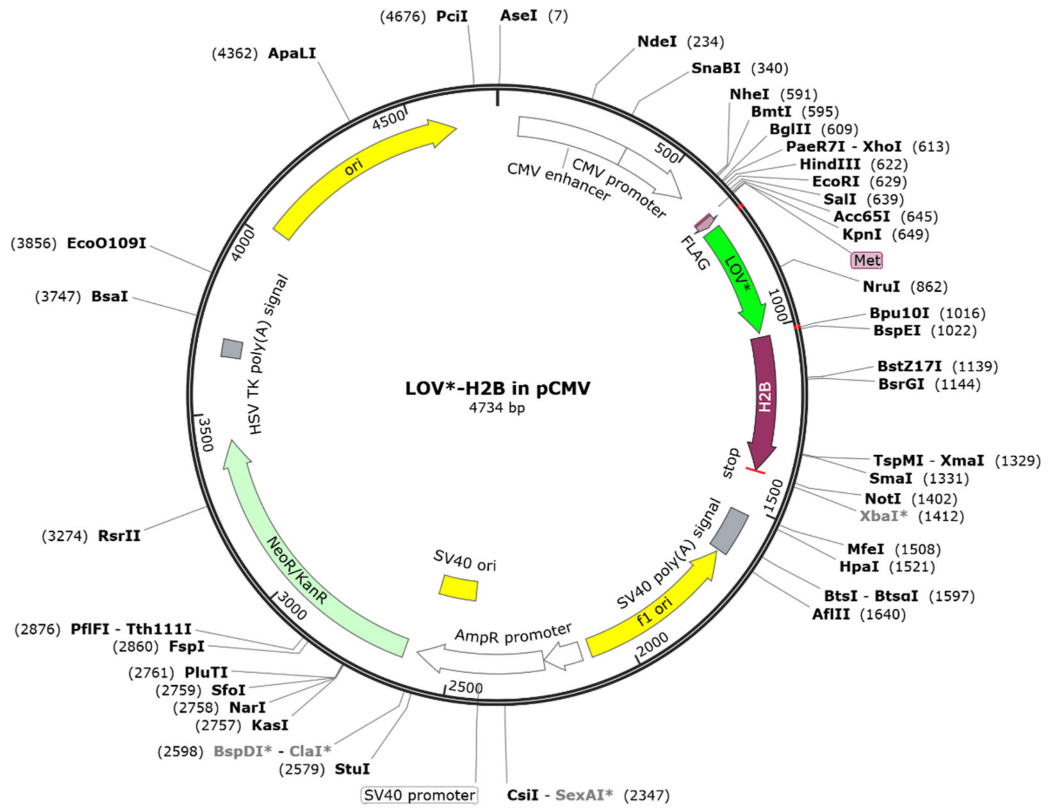


Expressed protein sequence:

MGDYKDDDDKGGSGMEKSFVITDPRLPDNPIIFASDGFLELTEYSREEILGRNGRFLQGPETDQATVQKIRDAIRDQREITVQLINYTKSGKKFWNLLHLQPMRDQKGELQYFIGVLLDGGGSGPEPAKSAPAPKKGSKKAVTKAQKKDGKKRKRSRKESYSVYVYKVLKQVHPDTGISSKAMGIMNSFVNDIFERIAGEASRLAHYNKRSTITSREIQTAVRLLLPGELAKHAVSEGTKAVTKYTSAK

### miniSOG-H2B in pCMV

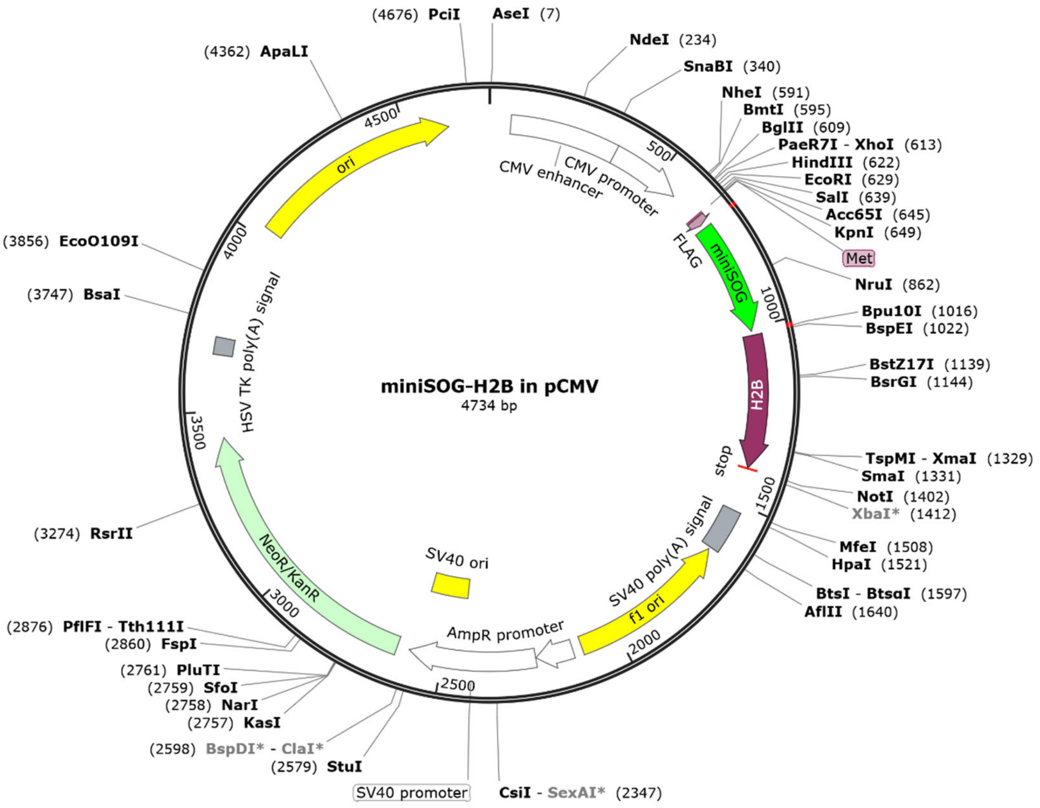


Expressed protein sequence:

MGDYKDDDDKGGSGMEKSFVITDPRLPDNPIIFASDGFLELTEYSREEILGRNGRFLQGPETDQATVQKIRDAIRDQREITVQLINYTKSGKKFWNLLHLQPMRDQKGELQYFIGVLLDGGGSGPEPAKSAPAPKKGSKKAVTKAQKKDGKKRKRSRKESYSVYVYKVLKQVHPDTGISSKAMGIMNSFVNDIFERIAGEASRLAHYNKRSTITSREIQTAVRLLLPGELAKHAVSEGTKAVTKYTSAK

miniSOG2-H2B in pCMV

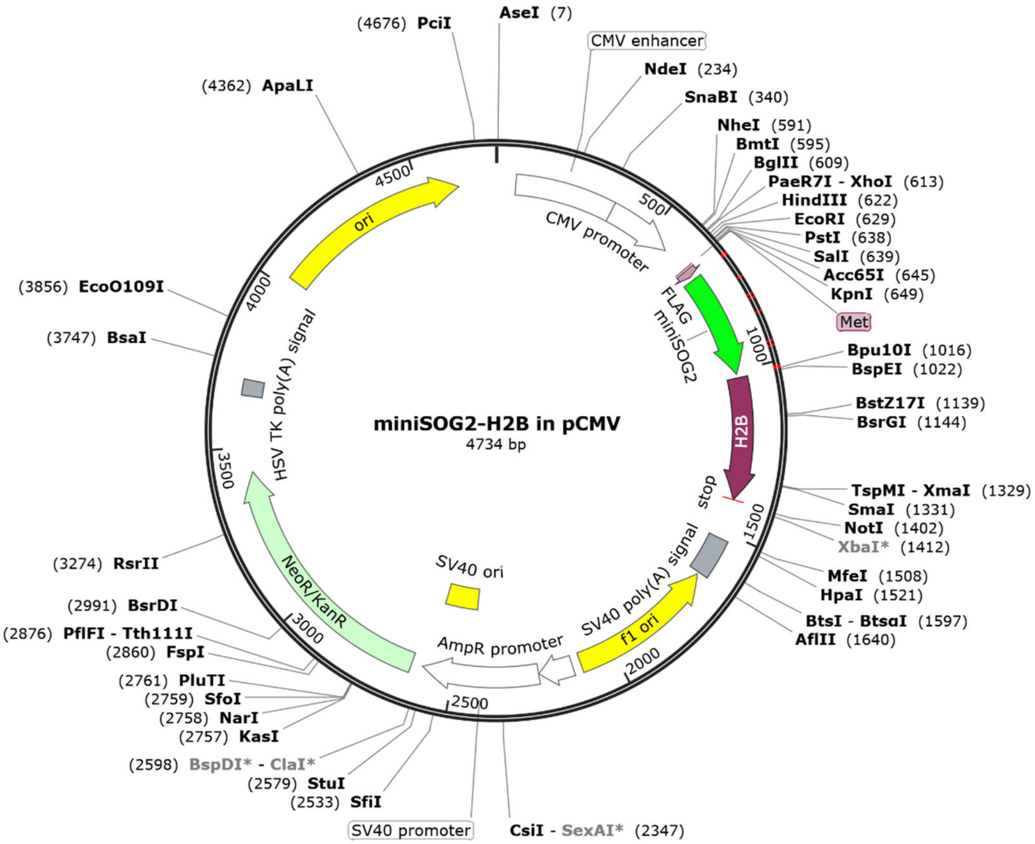


Expressed protein sequence:

MGDYKDDDDKGGSGMEKSFVITDPRLPDNPIIFASDSFLELTEYSREEILGRNPRFLRGPETDQATVQKIHDAIRDQREITVQLINYTKSGKKF
WNLFRLQPIRDQKGELQYFIGVQLDGGGSGPEPAKSAPAPKKGSKKAVTKAQKKDGKKRKRSRKESYSVYVYKVLKQVHPDTGISSKAMGI
MNSFVNDIFERIAGEASRLAHYNKRSTITSREIQTAVRLLLPGELAKHAVSEGTKAVTKYTSAK

<u>SOPP2-H2B in pCMV</u>

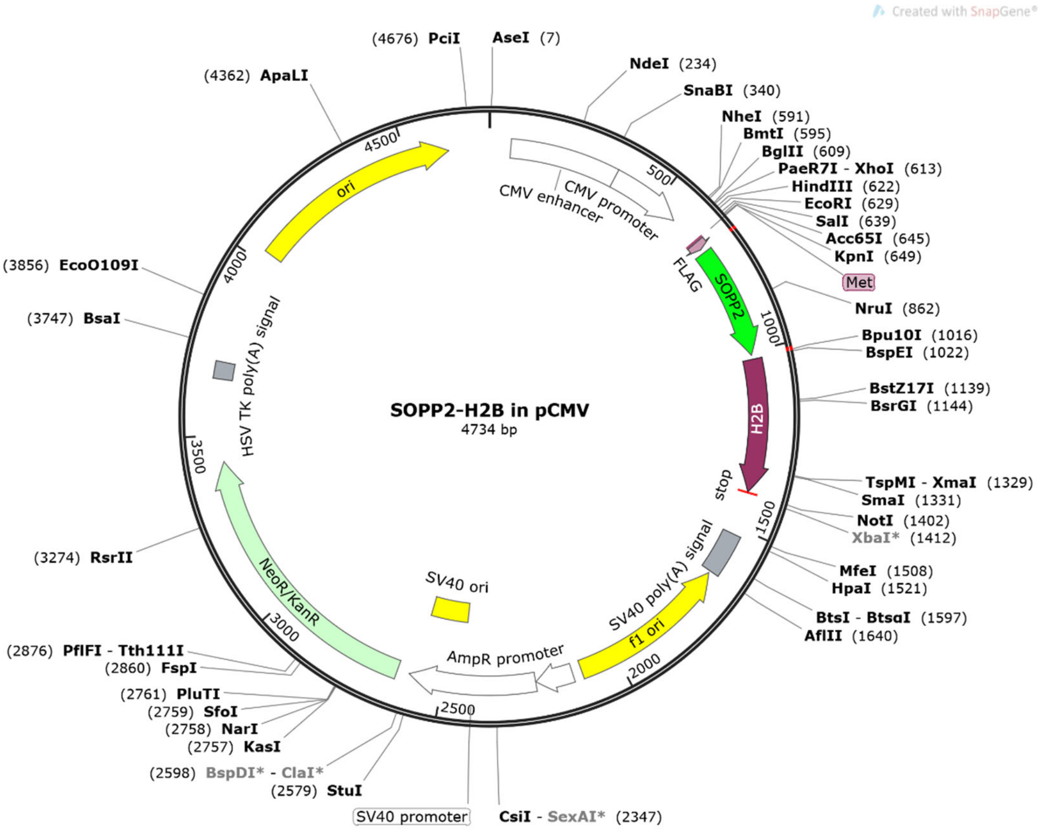


Expressed protein sequence:

MGDYKDDDDKGGSGMEKSFVITDPRLPDNPIIFASDGFLELTEYSREEILGRNGRFLQGPETDQATVQKIRDAIRDQREITVQLINYTKSGKK

FLNLLHLQPMRDQKGELQYFIGVVLDGGGSGPEPAKSAPAPKKGSKKAVTKAQKKDGKKRKRSRKESYSVYVYKVLKQVHPDTGISSKAM

GIMNSFVNDIFERIAGEASRLAHYNKRSTITSREIQTAVRLLLPGELAKHAVSEGTKAVTKYTSAK

SOPP3-H2B in pCMV

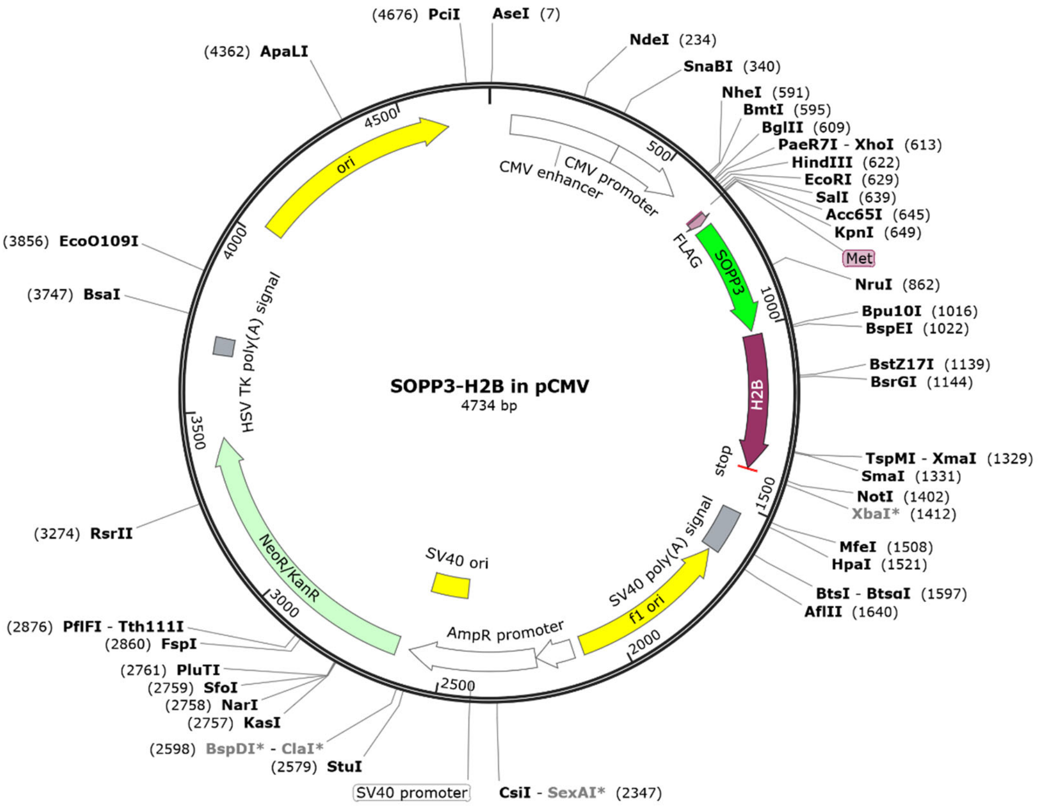


Expressed protein sequence:

MGDYKDDDDKGGSGMEKSFVITDPRLPDNPIIFASDGFLELTEYSREEILGRNGRFLQGPETDQATVQKIRDAIRDQREITVQLINYTKSGKK

FLNLLNLQPIRDQKGELQAFIGVVLDGGGSGPEPAKSAPAPKKGSKKAVTKAQKKDGKKRKRSRKESYSVYVYKVLKQVHPDTGISSKAMGI

MNSFVNDIFERIAGEASRLAHYNKRSTITSREIQTAVRLLLPGELAKHAVSEGTKAVTKYTSAK

<u>LOV* in pCMV</u>

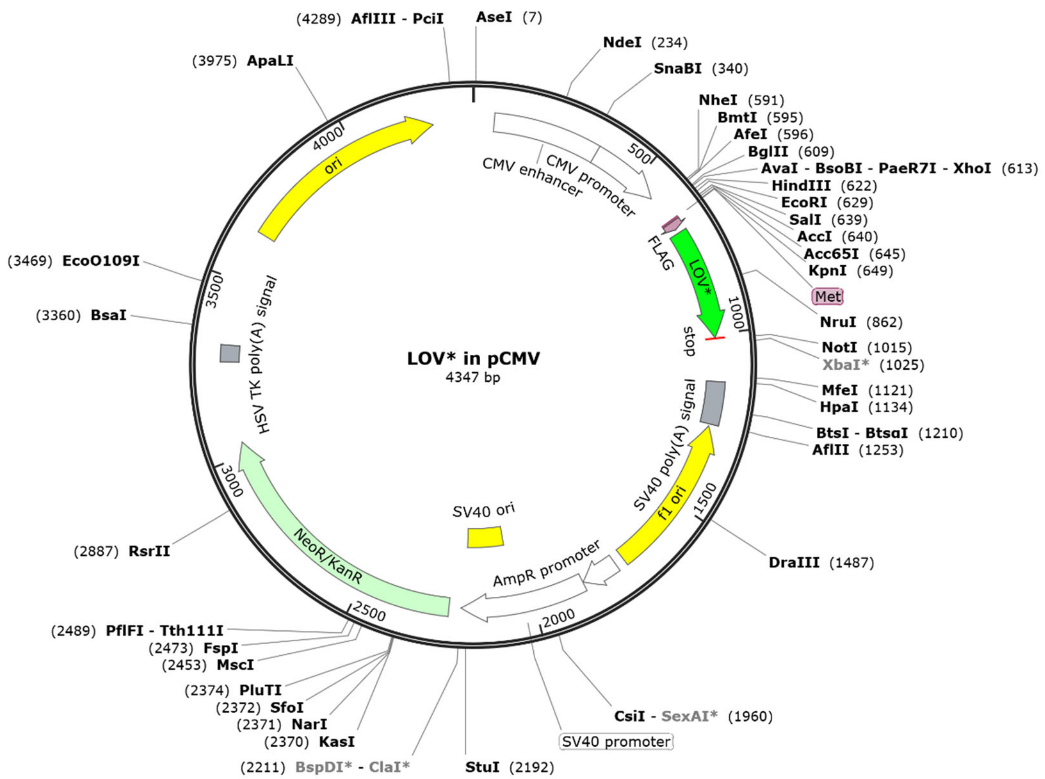


Expressed protein sequence:

MGDYKDDDDKGGSGMEKSFVITDPRLPDNPIIFASDGFLELTEYSREEILGRNGRFLQGPETDQATVQKIRDAIRDQREITVQLINYTKSGKK
FWNLLHLQPMRDQKGELQYFIGVLLDG

LOV*-LMNA in pCMV

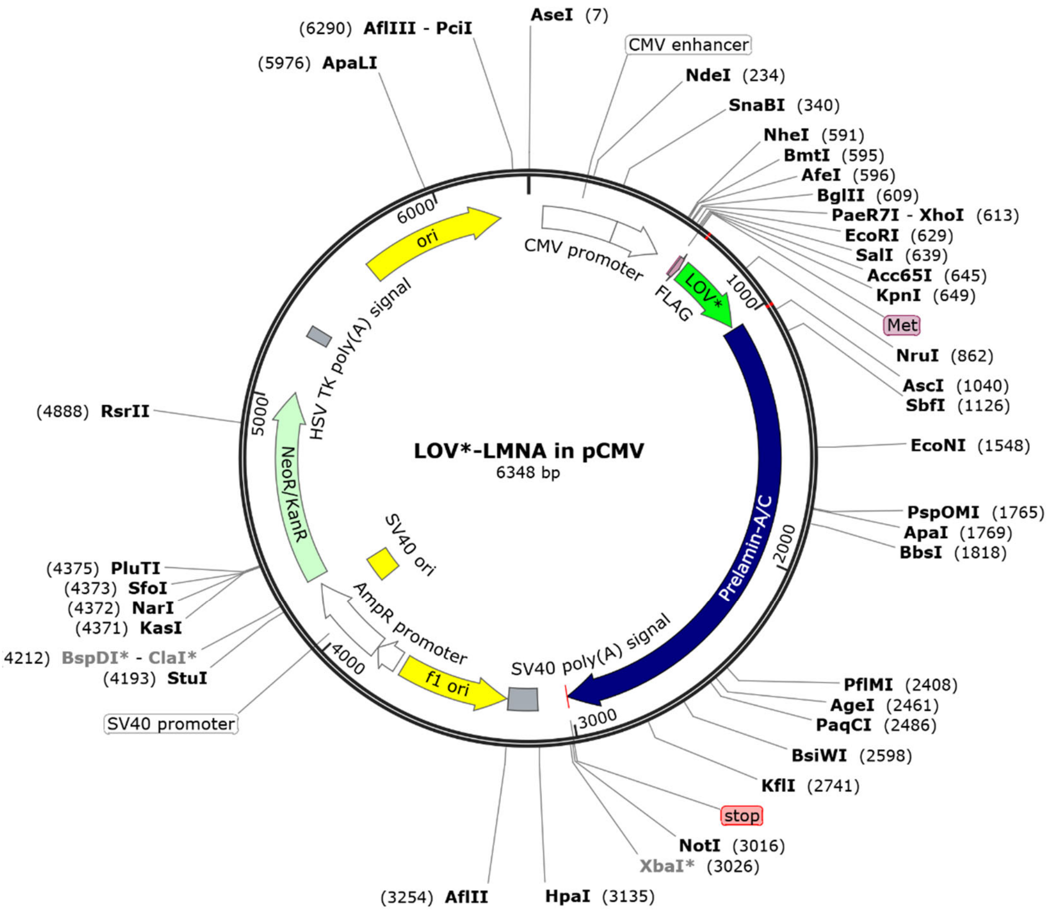


Expressed protein sequence:

MGDYKDDDDKGGSGMEKSFVITDPRLPDNPIIFASDGFLELTEYSREEILGRNGRFLQGPETDQATVQKIRDAIRDQREITVQLINYTKSGKK
FWNLLHLQPMRDQKGELQYFIGVLLDGGGSGETPSQRRATRSGAQASSTPLSPTRITRLQEKEDLQELNDRLAVYIDRVRSLETENAGLRLR
ITESEEVVSREVSGIKAAYEAELGDARKTLDSVAKERARLQLELSKVREEFKELKARNTKKEGDLIAAQARLKDLEALLNSKEAALSTALSEKRTL
EGELHDLRGQVAKLEAALGEAKKQLQDEMLRRVDAENRLQTMKEELDFQKNIYSEELRETKRRHETRLVEIDNGKQREFESRLADALQELR
AQHEDQVEQYKKELEKTYSAKLDNARQSAERNSNLVGAAHEELQQSRIRIDSLSAQLSQLQKQLAAKEAKLRDLEDSLARERDTSRRLLAEK
EREMAEMRARMQQQLDEYQELLDIKLALDMEIHAYRKLLEGEEERLRLSPSPTSQRSRGRASSHSSQTQGGGSVTKKRKLESTESRSSFSQ
HARTSGRVAVEEVDEEGKFVRLRNKSNEDQSMGNWQIKRQNGDDPLLTYRFPPKFTLKAGQVVTIWAAGAGATHSPPTDLVWKAQNT
WGCGNSLRTALINSTGEEVAMRKLVRSVTVVEDDEDEDGDDLLHHHHGSHCSSSGDPAEYNLRSRTVLCGTCGQPADKASASGSGAQVG
GPISSGSSASSVTVTRSYRSVGGSGGGSFGDNLVTRSYLLGNSSPRTQSPQNCSIM

<u>PDK1-LOV* in pCMV</u>

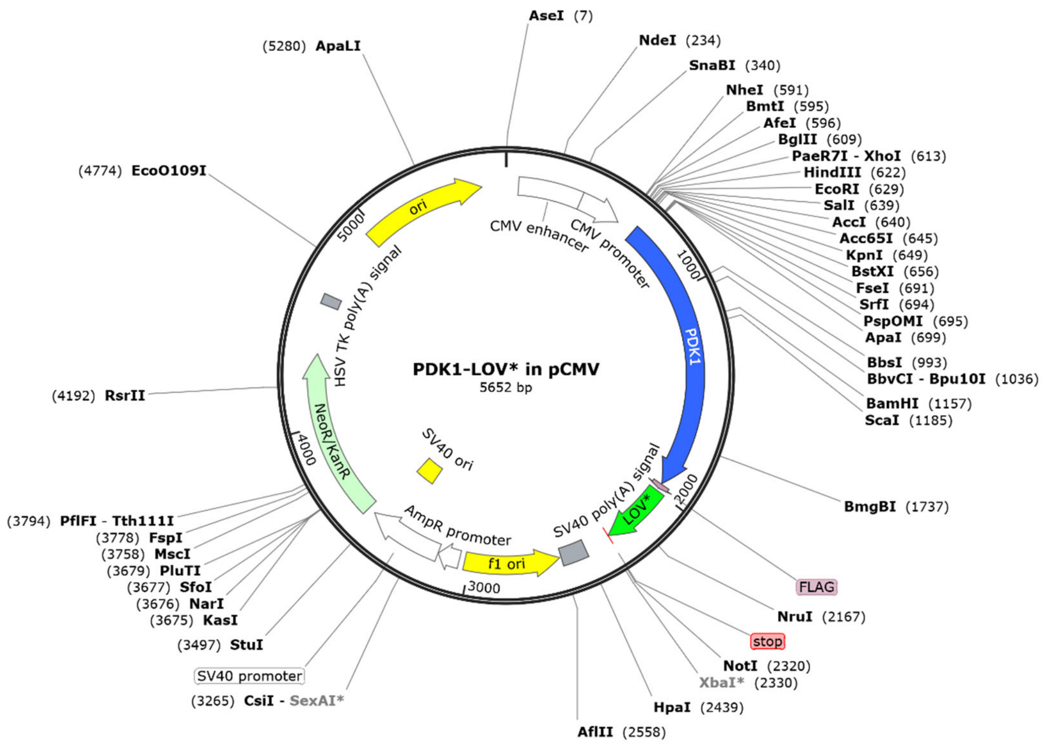


Expressed protein sequence:

MRLARLLRGAALAGPGPGLRAAGFSRSFSSDSGSSPASERGVPGQVDFYARFSPSPLSMKQFLDFGSVNACEKTSFMFLRQELPVRLANIM
KEISLLPDNLLRTPSVQLVQSWYIQSLQELLDFKDKSAEDAKAIYDFTDTVIRIRNRHNDVIPTMAQGVIEYKESFGVDPVTSQNVQYFLDRFY
MSRISIRMLLNQHSLLFGGKGKGSPSHRKHIGSINPNCNVLEVIKDGYENARRLCDLYYINSPELELEELNAKSPGQPIQVVYVPSHLYHMVF
ELFKNAMRATMEHHANRGVYPPIQVHVTLGNEDLTVKMSDRGGGVPLRKIDRLFNYMYSTAPRPRVETSRAVPLAGFGYGLPISRLYAQY
FQGDLKLYSLEGYGTDAVIYIKALSTDSIERLPVYNKAAWKHYNTNHEADDWCVPSREPKDMTTFRSAGDYKDDDDKGGSGMEKSFVITD
PRLPDNPIIFASDGFLELTEYSREEILGRNGRFLQGPETDQATVQKIRDAIRDQREITVQLINYTKSGKKFWNLLHLQPMRDQKGELQYFIGVL
LDG

<u>LOV*-3xNIK in pCMV</u>

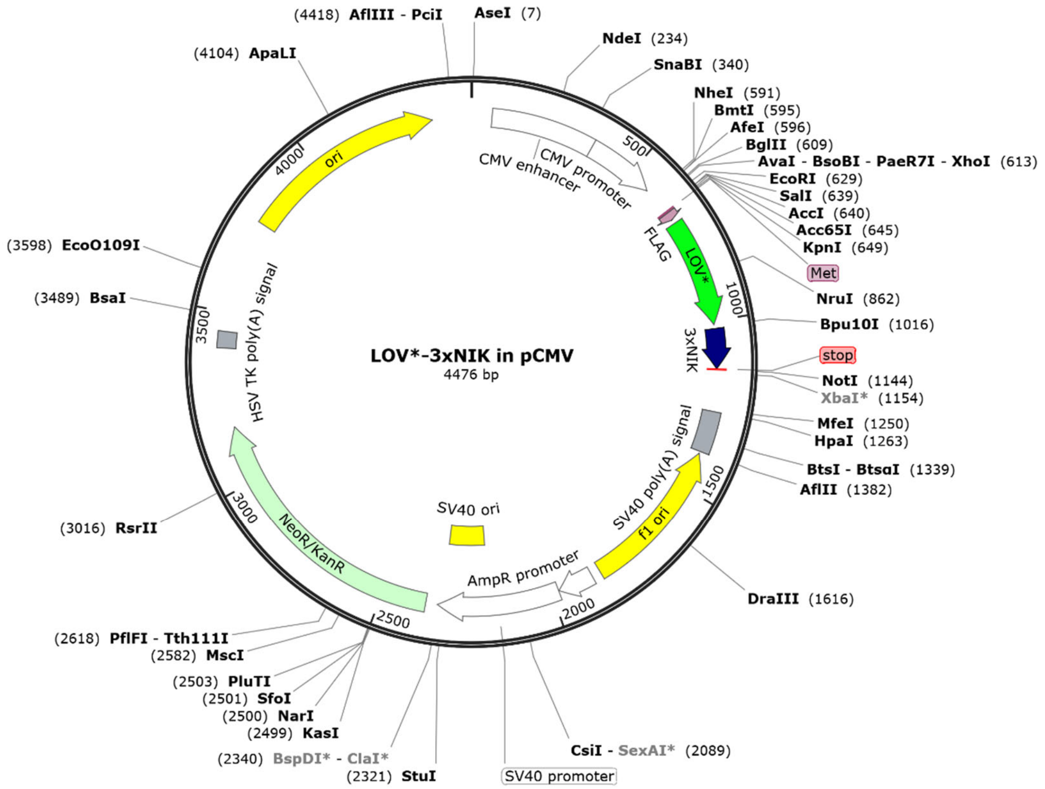


Expressed protein sequence:

MGDYKDDDDKGGSGMEKSFVITDPRLPDNPIIFASDGFLELTEYSREEILGRNGRFLQGPETDQATVQKIRDAIRDQREITVQLINYTKSGKK
FWNLLHLQPMRDQKGELQYFIGVLLDGGGSGRMRKKRKKKLRILRMRKKRKKKLRILRMRKKRKKKLRIL

LOV*-B7 in pCMV

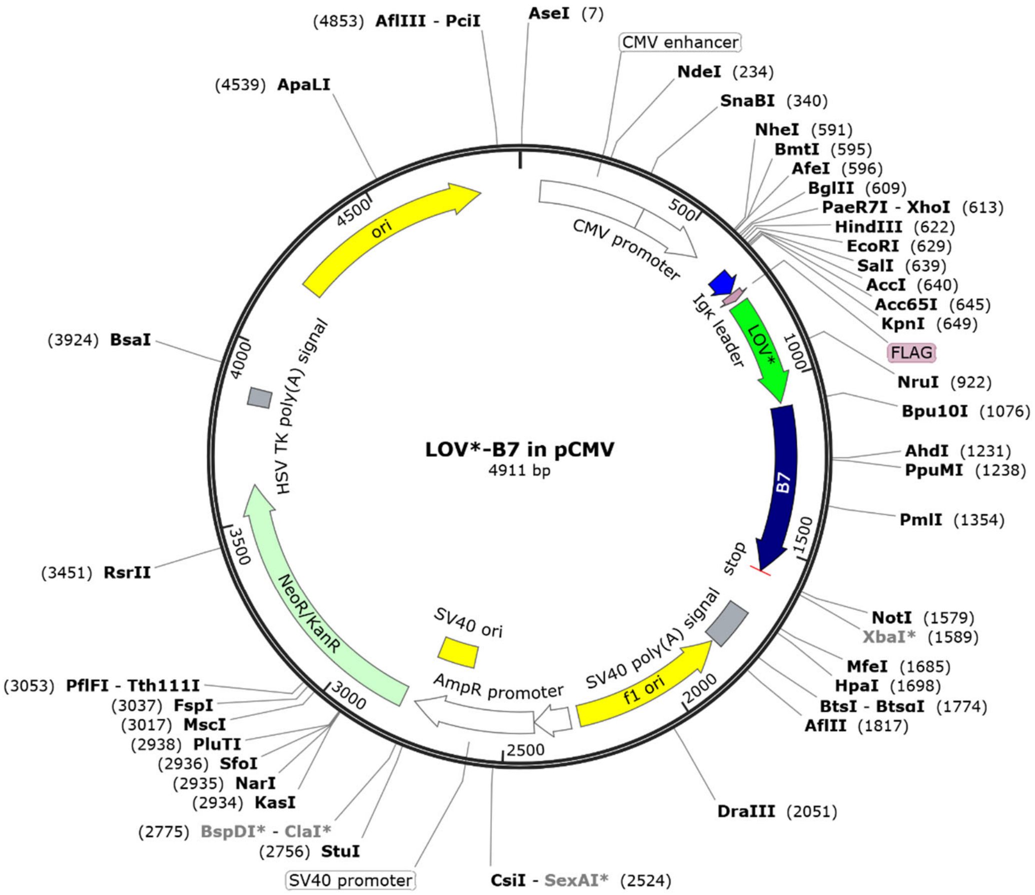


Expressed protein sequence:

METDTLLLWVLLLWVPGSTGDGDYKDDDDKGGSGMEKSFVITDPRLPDNPIIFASDGFLELTEYSREEILGRNGRFLQGPETDQATVQKIR
DAIRDQREITVQLINYTKSGKKFWNLLHLQPMRDQKGELQYFIGVLLDGGGSGADFSTPNITESGNPSADTKRITCFASGGFPKPRFSWLEN
GRELPGINTTISQDPESELYTISSQLDFNTTRNHTIKCLIKYGDAHVSEDFTWEKPPEDPPDSKNTLVLFGAGFGAVITVVVIVVIIKCFCKHRSC
FRRNEASRETNNSLTFGPEEALAEQTVFL

MTS-LOV* in pLenti-EF1a

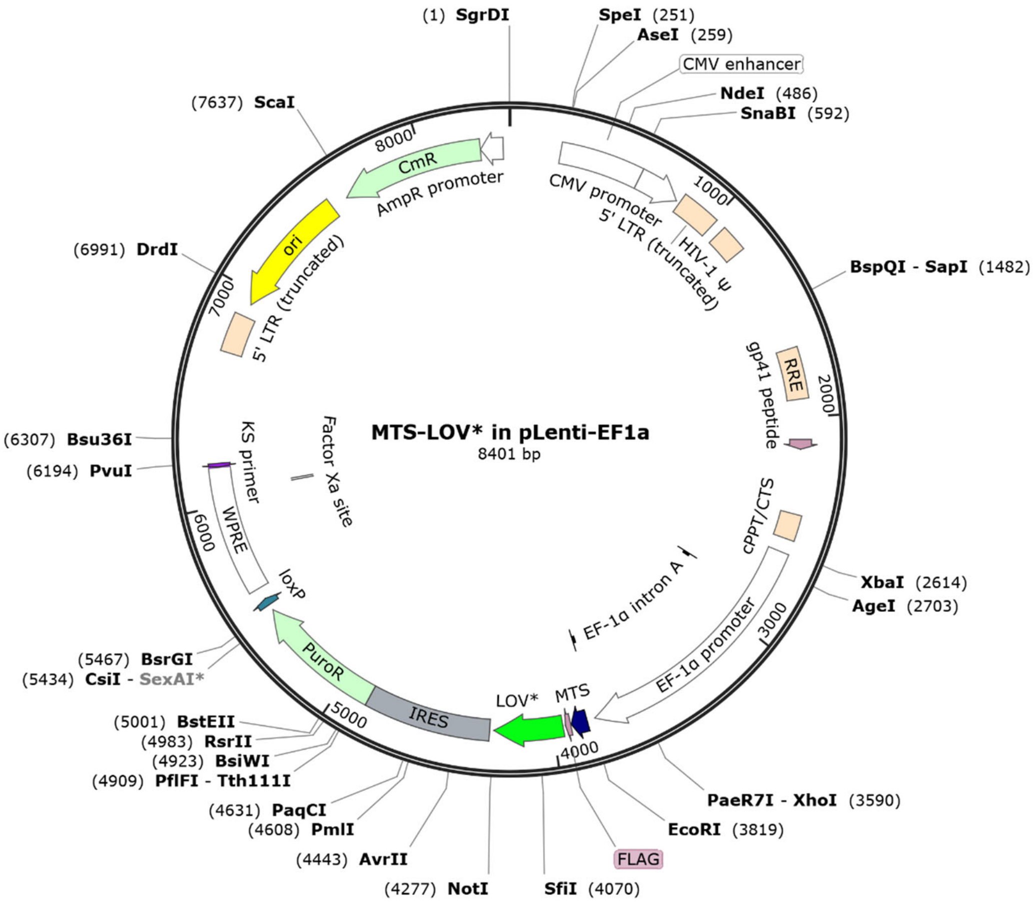


Expressed protein sequence:

MLATRVFSLVGKRAISTSVCVRAHKGDYKDDDDKGGSGMEKSFVITDPRLPDNPIIFASDGFLELTEYSREEILGRNGRFLQGPETDQATVQKIRDAIRDQREITVQLINYTKSGKKFWNLLHLQPMRDQKGELQYFIGVLLDG

PARP1-LOV* in PB-TRE3G

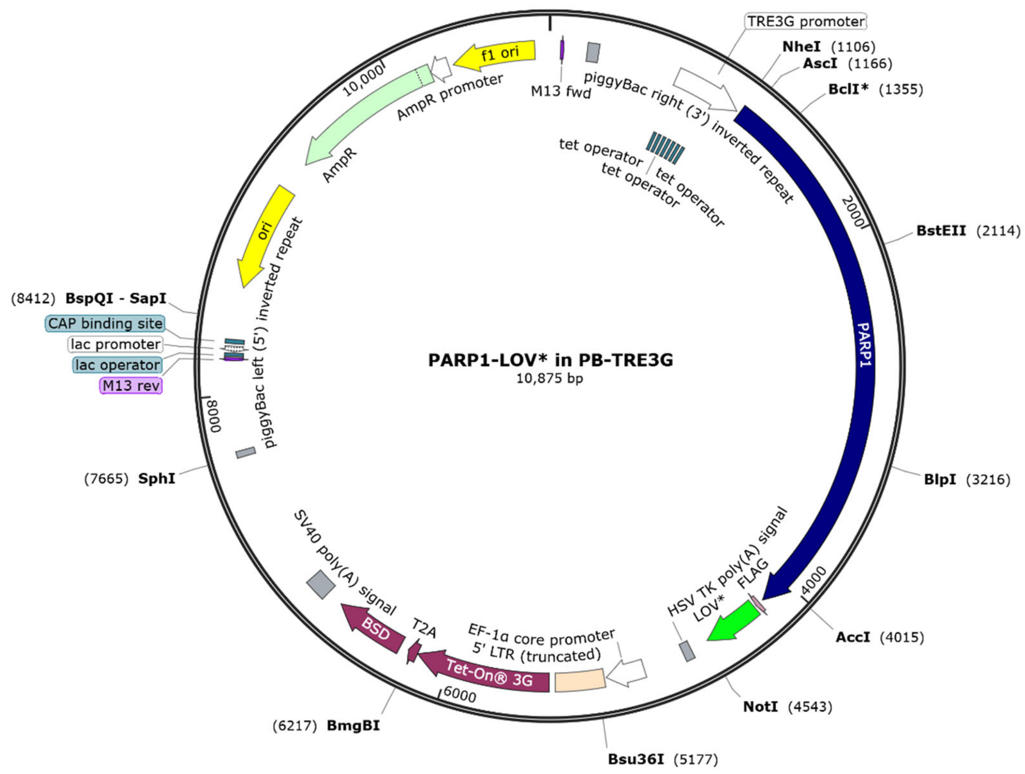


Expressed protein sequence:

MAESSDKLYRVEYAKSGRASCKKCSESIPKDSLRMAIMVQSPMFDGKVPHWYHFSCFWKVGHSIRHPDVEVDGFSELRWDDQQKVKKT
AEAGGVTGKGQDGIGSKAEKTLGDFAAEYAKSNRSTCKGCMEKIEKGQVRLSKKMVDPEKPQLGMIDRWYHPGCFVKNREELGFRPEYS
ASQLKGFSLLATEDKEALKKQLPGVKSEGKRKGDEVDGVDEVAKKKSKKEKDKDSKLEKALKAQNDLIWNIKDELKKVCSTNDLKELLIFNKQ
QVPSGESAILDRVADGMVFGALLPCEECSGQLVFKSDAYYCTGDVTAWTKCMVKTQTPNRKEWVTPKEFREISYLKKLKVKKQDRIFPPET
SASVAATPPPSTASAPAAVNSSASADKPLSNMKILTLGKLSRNKDEVKAMIEKLGGKLTGTANKASLCISTKKEVEKMNKKMEEVKEANIRV
VSEDFLQDVSASTKSLQELFLAHILSPWGAEVKAEPVEVVAPRGKSGAALSKKSKGQVKEEGINKSEKRMKLTLKGGAAVDPDSGLEHSAH
VLEKGGKVFSATLGLVDIVKGTNSYYKLQLLEDDKENRYWIFRSWGRVGTVIGSNKLEQMPSKEDAIEHFMKLYEEKTGNAWHSKNFTKYP
KKFYPLEIDYGQDEEAVKKLTVNPGTKSKLPKPVQDLIKMIFDVESMKKAMVEYEIDLQKMPLGKLSKRQIQAAYSILSEVQQAVSQGSSDS
QILDLSNRFYTLIPHDFGMKKPPLLNNADSVQAKAEMLDNLLDIEVAYSLLRGGSDDSSKDPIDVNYEKLKTDIKVVDRDSEEAEIIRKYVKNT
HATTHNAYDLEVIDIFKIEREGECQRYKPFKQLHNRRLLWHGSRTTNFAGILSQGLRIAPPEAPVTGYMFGKGIYFADMVSKSANYCHTSQG
DPIGLILLGEVALGNMYELKHASHISKLPKGKHSVKGLGKTTPDPSANISLDGVDVPLGTGISSGVNDTSLLYNEYIVYDIAQVNLKYLLKLKFN
FKTSLWGGSGGSMGDYKDDDDKGGSGMEKSFVITDPRLPDNPIIFASDGFLELTEYSREEILGRNGRFLQGPETDQATVQKIRDAIRDQREI
TVQLINYTKSGKKFWNLLHLQPMRDQKGELQYFIGVLLDG

LOV*-MVP in pCMV

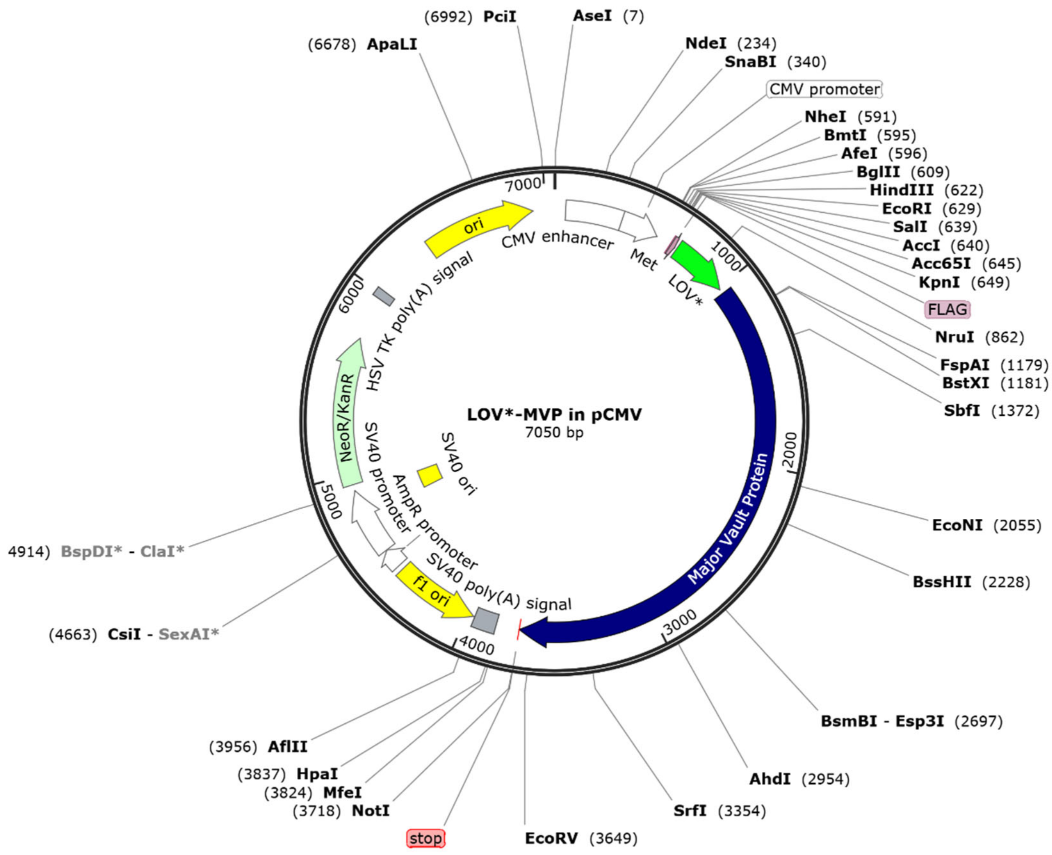


Expressed protein sequence:

MGDYKDDDDKGGSGMEKSFVITDPRLPDNPIIFASDGFLELTEYSREEILGRNGRFLQGPETDQATVQKIRDAIRDQREITVQLINYTKSGKKFWNLLHLQPMRDQKGELQYFIGVLLDGGGSGGGSGMATEEFIIRIPPYHYIHVLDQNSNVSRVEVGPKTYIRQDNERVLFAPMRMVTVPPRHYCTVANPVSRDAQGLVLFDVTGQVRLRHADLEIRLAQDPFPLYPGEVLEKDITPLQVVLPNTALHLKALLDFEDKDGDKVVAGDEWLFEGPGTYIPRKEVEVVEIIQATIIRQNQALRLRARKECWDRDGKERVTGEEWLVTTVGAYLPAVFEEVLDLVDAVILTEKTALHLRARRNFRDFRGVSRRTGEEWLVTVQDTEAHVPDVHEEVLGVVPITTLGPHNYCVILDPVGPDGKNQLGQKRVVKGEKSFFLQPGEQLEQGIQDVYVLSEQQGLLLRALQPLEEGEDEEKVSHQAGDHWLIRGPLEYVPSAKVEVVEERQAIPLDENEGIYVQDVKTGKVRAVIGSTYMLTQDEVLWEKELPPGVEELLNKGQDPLADRGEKDTAKSLQPLAPRNKTRVVSYRVPHNAAVQVYDYREKRARVVFGPELVSLGPEEQFTVLSLSAGRPKRPHARRALCLLLGPDFFTDVITIETADHARLQLQLAYNWHFEVNDRKDPQETAKLFSVPDFVGDACKAIASRVRGAVASVTFDDFHKNSARIIRTAVFGFETSEAKGPDGMALPRPRDQAVFPQNGLVVSSVDVQSVEPVDQRTRDALQRSVQLAIEITTNSQEAAAKHEAQRLEQEARGRLERQKILDQSEAEKARKELLELEALSMAVESTGTAKAEAESRAEAARIEGEGSVLQAKLKAQALAIETEAELQRVQKVRELELVYARAQLELEVSKAQQLAEVEVKKFKQMTEAIGPSTIRDLAVAGPEMQVKLLQSLGLKSTLITDGSTPINLFNTAFGLLGMGPEGQPLGRRVASGPSPGEGISPQSAQAPQAPGDNHVVPVLR

<u>MVP-LOV* in pCMV</u>

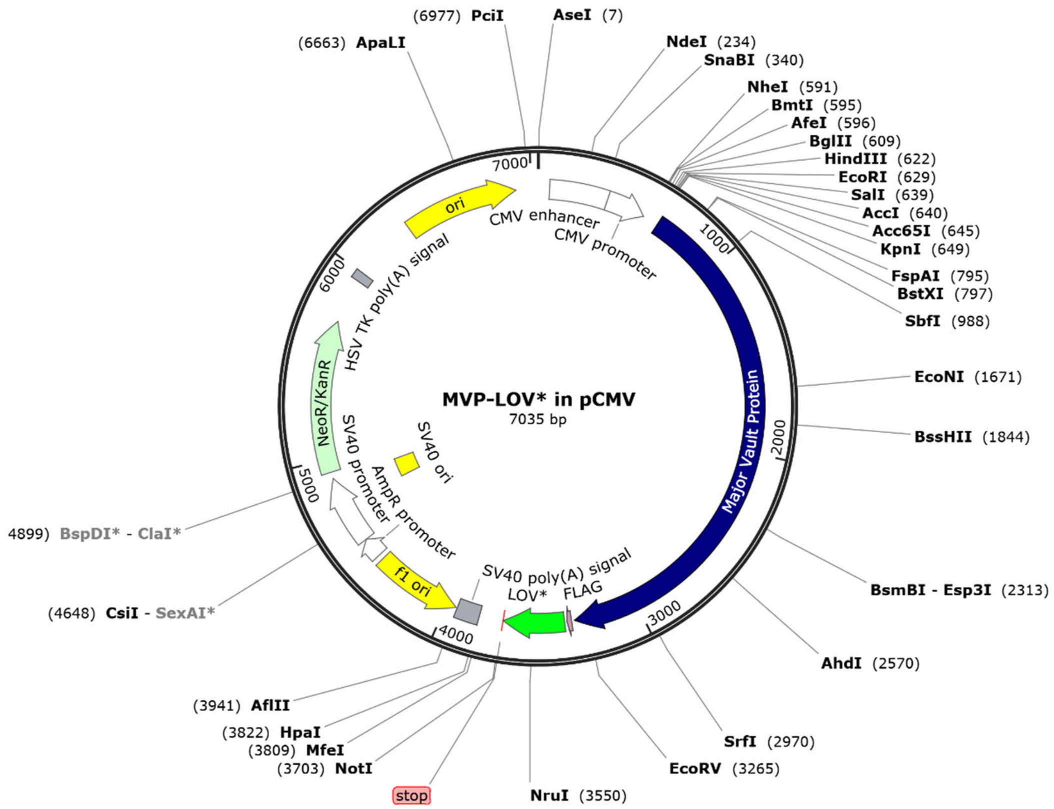


Expressed protein sequence:

MATEEFIIRIPPYHYIHVLDQNSNVSRVEVGPKTYIRQDNERVLFAPMRMVTVPPRHYCTVANPVSRDAQGLVLFDVTGQVRLRHADLEIR
LAQDPFPLYPGEVLEKDITPLQVVLPNTALHLKALLDFEDKDGDKVVAGDEWLFEGPGTYIPRKEVEVVEIIQATIIRQNQALRLRARKECWD
RDGKERVTGEEWLVTTVGAYLPAVFEEVLDLVDAVILTEKTALHLRARRNFRDFRGVSRRTGEEWLVTVQDTEAHVPDVHEEVLGVVPITT
LGPHNYCVILDPVGPDGKNQLGQKRVVKGEKSFFLQPGEQLEQGIQDVYVLSEQQGLLLRALQPLEEGEDEEKVSHQAGDHWLIRGPLEY
VPSAKVEVVEERQAIPLDENEGIYVQDVKTGKVRAVIGSTYMLTQDEVLWEKELPPGVEELLNKGQDPLADRGEKDTAKSLQPLAPRNKTR
VVSYRVPHNAAVQVYDYREKRARVVFGPELVSLGPEEQFTVLSLSAGRPKRPHARRALCLLLGPDFFTDVITIETADHARLQLQLAYNWHFE
VNDRKDPQETAKLFSVPDFVGDACKAIASRVRGAVASVTFDDFHKNSARIIRTAVFGFETSEAKGPDGMALPRPRDQAVFPQNGLVVSSV
DVQSVEPVDQRTRDALQRSVQLAIEITTNSQEAAAKHEAQRLEQEARGRLERQKILDQSEAEKARKELLELEALSMAVESTGTAKAEAESRA
EAARIEGEGSVLQAKLKAQALAIETEAELQRVQKVRELELVYARAQLELEVSKAQQLAEVEVKKFKQMTEAIGPSTIRDLAVAGPEMQVKLL
QSLGLKSTLITDGSTPINLFNTAFGLLGMGPEGQPLGRRVASGPSPGEGISPQSAQAPQAPGDNHVVPVLRGGSGGDYKDDDDKGGSGM
EKSFVITDPRLPDNPIIFASDGFLELTEYSREEILGRNGRFLQGPETDQATVQKIRDAIRDQREITVQLINYTKSGKKFWNLLHLQPMRDQKGE
LQYFIGVLLDG

LOV* in pET30

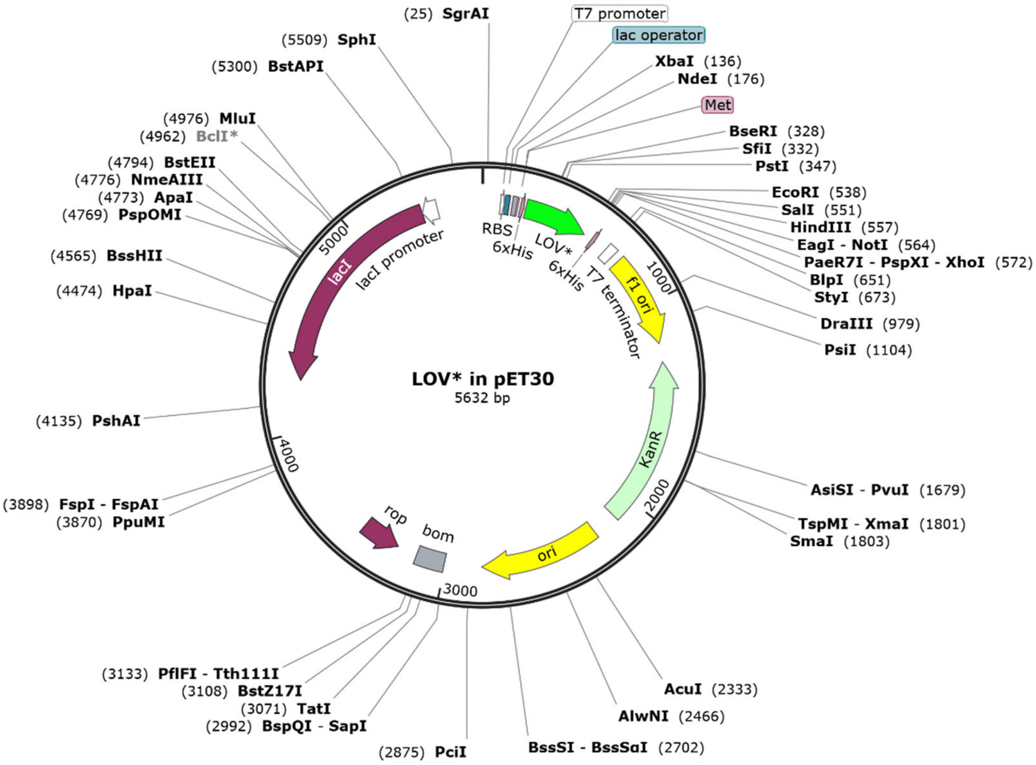


Expressed protein sequence:

MGSSHHHHHHGSGMEKSFVITDPRLPDNPIIFASDGFLELTEYSREEILGRNGRFLQGPETDQATVQKIRDAIRDQREITVQLINYTKSGKKF
WNLLHLQPMRDQKGELQYFIGVLLDG

<u>eGFP-HMGB1 in pcDNA3.1</u>

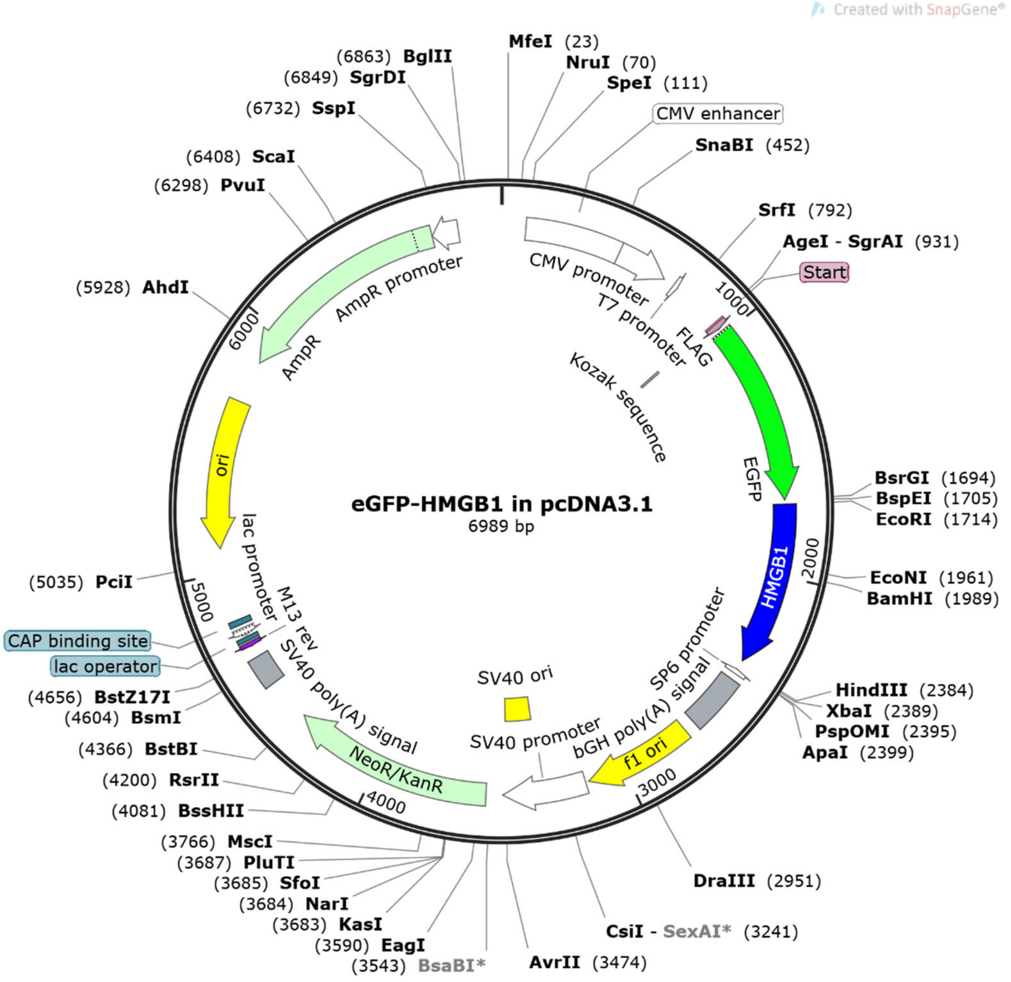


Expressed protein sequence:

MGDYKDDDDKGGSGMVSKGEELFTGVVPILVELDGDVNGHKFSVSGEGEGDATYGKLTLKFICTTGKLPVPWPTLVTTLTYGVQCFSRYP
DHMKQHDFFKSAMPEGYVQERTIFFKDDGNYKTRAEVKFEGDTLVNRIELKGIDFKEDGNILGHKLEYNYNSHNVYIMADKQKNGIKVNF
KIRHNIEDGSVQLADHYQQNTPIGDGPVLLPDNHYLSTQSALSKDPNEKRDHMVLLEFVTAAGITLGMDELYKGSGSEFMGKGDPKKPRG
KMSSYAFFVQTCREEHKKKHPDASVNFSEFSKKCSERWKTMSAKEKGKFEDMAKADKARYEREMKTYIPPKGETKKKFKDPNAPKRPPSA
FFLFCSEYRPKIKGEHPGLSIGDVAKKLGEMWNNTAADDKQPYEKKAAKLKEKYEKDIAAYRAKGKPDAAKKGVVKAEKSKKKKEEEEDEED
EEDEEEEEDEEDEDEEEDDDDE

**Recombinant 6xHis-LOV*:**

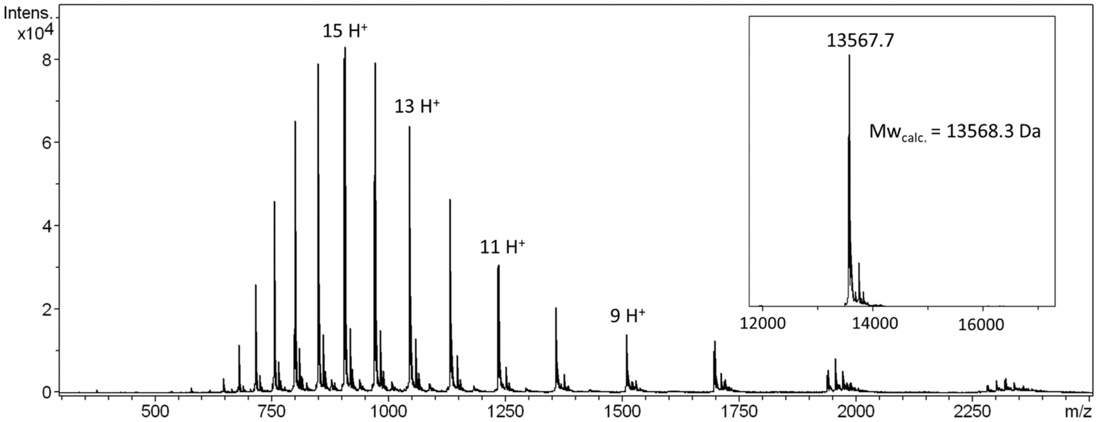


ESI-MS characterization of the recombinantly-expressed LOV* protein. Inset, deconvoluted MS.

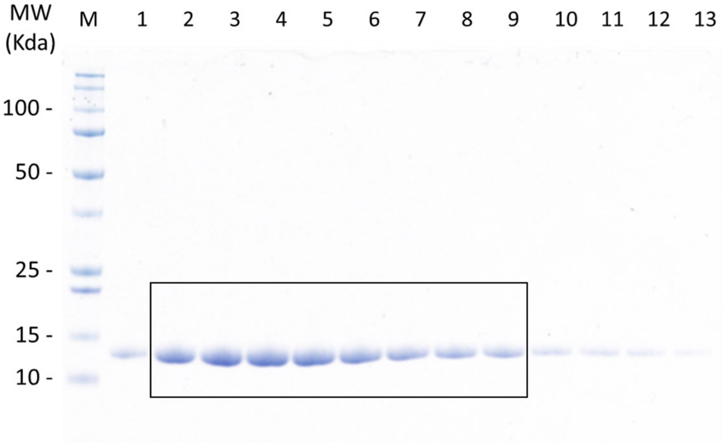


SDS-PAGE of the fractions of the major protein peak in the LOV* gel filtration chromatogram. Proteins were stained by Coomassie Blue. Fractions 2 to 9 were pooled and stored for further use.

### References


1. Shu, X. et al. A Genetically Encoded Tag for Correlated Light and Electron Microscopy of Intact Cells, Tissues, and Organisms. *PLOS Biology* **9**, e1001041 (2011).
2. Westberg, M., Holmegaard, L., Pimenta, F.M., Etzerodt, M. & Ogilby, P.R. Rational Design of an Efficient, Genetically Encodable, Protein-Encased Singlet Oxygen Photosensitizer. *Journal of the American Chemical Society* **137**, 1632-1642 (2015).
3. Westberg, M., Bregnhøj, M., Etzerodt, M. & Ogilby, P.R. No Photon Wasted: An Efficient and Selective Singlet Oxygen Photosensitizing Protein. *The Journal of Physical Chemistry B* **121**, 9366-9371 (2017).
4. Makhijani, K. et al. Precision Optogenetic Tool for Selective Single- and Multiple-Cell Ablation in a Live Animal Model System. *Cell Chemical Biology* **24**, 110-119 (2017).
5. Bissonnette, N.B. et al. Design of a Multiuse Photoreactor To Enable Visible-Light Photocatalytic Chemical Transformations and Labeling in Live Cells. *ChemBioChem* **21**, 3555-3562 (2020).
6. Suatoni, J.C., Snyder, R.E. & Clark, R.O. Voltammetric studies of phenol and aniline ring substitution. *Analytical Chemistry* **33**, 1894-1897 (1961).
7. Yoshimura, A. & Ohno, T. Lumiflavin-Sensitized Photooxygenation of Indole. *Photochemistry and Photobiology* **48**, 561-565 (1988).
8. Tay, N.E. et al. Targeted Activation in Localized Protein Environments via Deep Red Photoredox Catalysis. *ChemRxiv* 10.26434/chemrxiv-2021-x9bjv (2021).
9. Geri Jacob, B. et al. Microenvironment mapping via Dexter energy transfer on immune cells. *Science* **367**, 1091-1097 (2020).
10. Nguyen, U.T.T. et al. Accelerated chromatin biochemistry using DNA-barcoded nucleosome libraries. *Nature Methods* **11**, 834-840 (2014).
11. Birbach, A., Bailey, S.T., Ghosh, S. & Schmid, J.A. Cytosolic, nuclear and nucleolar localization signals determine subcellular distribution and activity of the NF-κB inducing kinase NIK. *Journal of Cell Science* **117**, 3615-3624 (2004).
12. Chen, K.C. et al. Membrane-localized activation of glucuronide prodrugs by β-glucuronidase enzymes. *Cancer Gene Therapy* **14**, 187-200 (2007).
13. Liszczak, G., Diehl, K.L., Dann, G.P. & Muir, T.W. Acetylation blocks DNA damage–induced chromatin ADP-ribosylation. *Nature Chemical Biology* **14**, 837-840 (2018).
14. Cox, J. & Mann, M. MaxQuant enables high peptide identification rates, individualized p.p.b.-range mass accuracies and proteome-wide protein quantification. *Nature Biotechnology* **26**, 1367-1372 (2008).
15. Hung, V. et al. Spatially resolved proteomic mapping in living cells with the engineered peroxidase APEX2. *Nature Protocols* **11**, 456-475 (2016).
16. Fazal, F.M. et al. Atlas of Subcellular RNA Localization Revealed by APEX-Seq. *Cell* **178**, 473-490.e26 (2019).
